# Feasibility of an eDNA-based educational program for high school students

**DOI:** 10.1101/2025.09.02.672770

**Authors:** Ryota P. Kitani, Tatsuya Saga, Minoru Kasada, Mieko Kiyono, Masayuki Sato, Atushi Ushimaru, Toshifumi Minamoto

## Abstract

1. Environmental education plays a crucial role in raising public awareness of ecosystem conservation. Environmental DNA (eDNA) analysis is a promising environmental education tool because it enables easy and comprehensive biodiversity assessment through simple water sampling. Despite several citizen-based surveys using eDNA analysis, minimal attention has been paid to its educational effectiveness.
2. In this study, we examined the educational feasibility of eDNA analysis by conducting an environmental education program that incorporated eDNA-based local fish surveys in three high schools in Japan. Through detailed questionnaire surveys conducted before and after the program, we verified whether our program increased student interest in biodiversity and ecosystem services.
3. In the program, students proactively identified diverse fish species living in the survey area by simply collecting water. The questionnaire survey results showed that student knowledge of and interest in biodiversity and ecosystem services increased after the program. Students became aware of local fish diversity and expressed their motivation to engage in conservation activities. We confirmed the educational impact of the program not only at the end but also immediately following the water sampling experience.
4. *Synthesis and applications.* These results provide the first empirical evidence of the positive effects of eDNA analysis on student environmental awareness and engagement. Our program provides a feasible model for eDNA-based environmental education. Because eDNA analysis can be applied to diverse target taxa that are unique to a given area, similar environmental education programs can be conducted for various citizen groups worldwide. Our research also suggested that eDNA-based environmental education was effective for younger generations with varying levels of interest and experience in nature, showing significant potential for addressing the extinction of natural experiences and motivating citizens to participate in ecosystem conservation activities.

## Introduction

Since the 20^th^ century, biodiversity has been declining globally (Finn et al., 2023), causing loss or deterioration of ecosystem services which are closely linked to biodiversity (Pereira et al., 2024). Ecosystems worldwide have been negatively affected by various human impacts, such as: land-use changes; global climate change, and; soil, water, and air eutrophication and/or pollution (Sala et al., 2000; Hysa et al., 2024). Thus, each of these ecosystems is degraded by various factors, including effects that can lead to biodiversity loss. Therefore, it is necessary to implement biodiversity conservation suitable for each unique specific local ecosystem. To address this challenge, it is essential to obtain consent and support from citizens for biodiversity conservation. However, interest in and knowledge of local biodiversity is often limited (Santana et al., 2023), and interest in and attitudes towards nature are declining (Soga & Gaston, 2016). In addition, people who do not interact with nature in their youth prefer ecosystems with lower biodiversity and have more negative attitudes towards ecosystem conservation in adulthood (Sato et al., 2017; Aoshima et al., 2023). Therefore, citizens are likely to have less interest in biodiversity conservation. To promote local biodiversity conservation, it is necessary to develop effective environmental education approaches that improve citizen interest, awareness, and valuation of biodiversity and ecosystem services, and encourage conservation action.

Environmental education is usually effective in facilitating citizen awareness of biodiversity and can promote sustainable conservation behaviour among citizens (Monroe et al., 2008). Because the experience of investigating local organisms is thought to raise awareness of biodiversity, biological surveys have been conducted as environmental education tools (Fougere, 1998; Giusti, 2019). However, there are several limitations to these biological surveys, such as the difficulty of identifying species owing to the lack of experts (Hooykaas et al., 2019; Cavadino et al., 2024), and the lack of time available to build the capacity of citizens to identify species. Additionally, discovering or capturing organisms is often not straightforward and largely depends on individual skills (Diogo et al., 2017). Moreover, direct biological surveys may have negative effects on people who are uncomfortable with living organisms. To compensate for such limitations, novel environmental education tools that can identify multiple species simultaneously are applicable to diverse locations and easily accessible to people with varying levels of comfort with direct wildlife interaction.

Environmental DNA (eDNA) analysis is a potential tool for environmental education. eDNA refers to all DNA present in the environment, including intra- and extra-organismal DNA from microscopic and macroscopic organisms (Minamoto, 2022). eDNA analysis can be used to comprehensively detect many species without prior morphological knowledge. Several studies have comprehensively detected diversity of fish (Miya et al., 2015; Michael et al., 2019; Wu et al., 2023), mammals (Ushio et al., 2017), birds (Ushio et al., 2018), and others (Coghlan et al., 2019; Sakata et al., 2022; Takenaka et al., 2023). Furthermore, with technical support from researchers or specialised companies, eDNA-based biological surveys can be implemented by non-expert citizens because the only task required in the field is to collect environmental water.

In several studies, water sampling was conducted by citizens through citizen science (Biggs et al., 2015; Sune et al., 2022; Kondoh et al., 2024). Previous studies have mainly focused on the efficiency of simultaneous sampling with many citizens, and few have focused on environmental education for participants. Suzuki‒Ohno et al. (2023) showed that citizens deepened their understanding of biodiversity by participating in sampling; however, the extent to which participant interest in biodiversity changed during eDNA-based surveys remains unclear. In addition, as these activities are often conducted by volunteers, participants may be biased towards nature lovers. Therefore, it is necessary to test whether activities using eDNA are effective for people with various levels of nature relatedness. There are also a few examples of the use of eDNA analysis in school educationare no reports confirming its effectiveness when delivered during school hours. Extracurricular environmental education can provide learning opportunities (Abdela, 2022); however, student busyness and a lack of preparation can hinder this activity (Solehah et al., 2022). Therefore, it is important to incorporate eDNA-based educational programs into school lessons and measure their effectiveness in promoting student interest in and knowledge of biodiversity and ecosystem conservation.

This study investigated whether eDNA-based surveys of local biodiversity are a feasible environmental education tool. We targeted high school students, because in Japan, these students learn the importance of conserving biodiversity and ecosystems through observation and experimentation (Ministry of Education, Culture, Sports, Science and Technology, 2018). In this study, an environmental education program using eDNA analysis was implemented during class time in three Japanese high schools, and questionnaire surveys were conducted before and after the program to evaluate its feasibility.

## 2. Materials and methods

### 2-1. High schools where survey was conducted

This study implemented an education program using an eDNA-based fish survey for a wide range of students enrolled in various courses in three schools: Hidakamioka and Ena high schools in Gifu Prefecture, and Yaizuchuo High School in Shizuoka Prefecture (Table 1). At Hidakamioka High School, the program was implemented for all first-year students in a regular course; whereas, at Ena High School, it was conducted for all second-year students in a science and mathematics course (a curriculum specialising in mathematics and science). At Yaizuchuo High School, the program was implemented for all first-year students, second- and third-year students in science biology-focused classes (specialised biology) of the regular course, and second-year students in the liberal arts-focused classes of the regular course. Generally, Japanese high schools offer regular and/or special courses, and students choose from multiple specialised classes (e.g., science or liberal arts) when advancing to the next grade.

**Table 1.** Number of students who responded to each questionnaire.

| school | Grade and<br>Course of<br>students*1 | Implementation<br>date | Number of<br>responses<br>to QS1 | Number of<br>responses<br>to QS2 | Number of<br>responses<br>to QS3 | Number of<br>students who<br>answered all<br>questions |
| --- | --- | --- | --- | --- | --- | --- |
| Hidakamioka | first-year,<br>Regular Course | 14 June and<br>5 July 2023 | 33 | 37 | 32 | 26 |
| Ena | second-year,<br>Science and<br>Mathematics<br>course | 15 June and<br>12 July 2023 | 77 | 76 | 79 | 64 |
| Yaizuchuo<br>(done water<br>sampling) | first-year,<br>Regular Course | 26-28<br>September and<br>8-10 November<br>2023 | 261 | 270 | 247 | 200 |
| Yaizuchuo<br>(no water<br>sampling) | second and<br>third year,<br>Regular course | 26-28<br>September and<br>8-10 November<br>2023 | 180 | 182 | 183 | 136 |

\*1 In Japan, students generally spend 3 years in high school, beginning at 15 years-of-age. Typically, students progress to more specialised courses in their
second year. Japanese high schools generally offer regular and special courses.

### 2-2. Consideration and implementation of environmental education

The eDNA-based educational program was designed to span three class periods within a regular school timetable (Fig. 1). In this study, each class period ranged from 45‒50 min (note: typical high school class periods are < 60 min in Japan). The programme was divided into two sessions. The first session consisted of two classes and the second session consisted of one class. In the first session, the first and second classes were devoted to a preliminary explanation and water sampling at a river near a high school, respectively. The preliminary explanation covered basic ecological concepts, including biodiversity, ecosystem services, and services, as well as an introduction to eDNA and water sampling protocols. In the second session, after briefly reviewing the first piece of content, students were presented with the fish metabarcoding analysis results from their collected samples. This was followed by group discussions on biodiversity and ecosystem services in the local river. During this session, the students used their tablet devices to search for information regarding fish in which they were interested. To better understand the fish communities in the region, we provided fish community data obtained through eDNA analysis (unpublished data from Kitani and ANEMONE DB: 2021KibanSRUN01_20210615T0440-URF-Butokamabetsugawa_MiFish). Students then compared their results with those from other regions. To verify the feasibility of the program, questionnaires were administered to the students three times: QS1 (before the activity), QS2 (after water sampling), and QS3 (after the entire lesson) (hereafter, QS1, QS2, and QS3 refer to individual questionnaires).

**Fig. 1.**
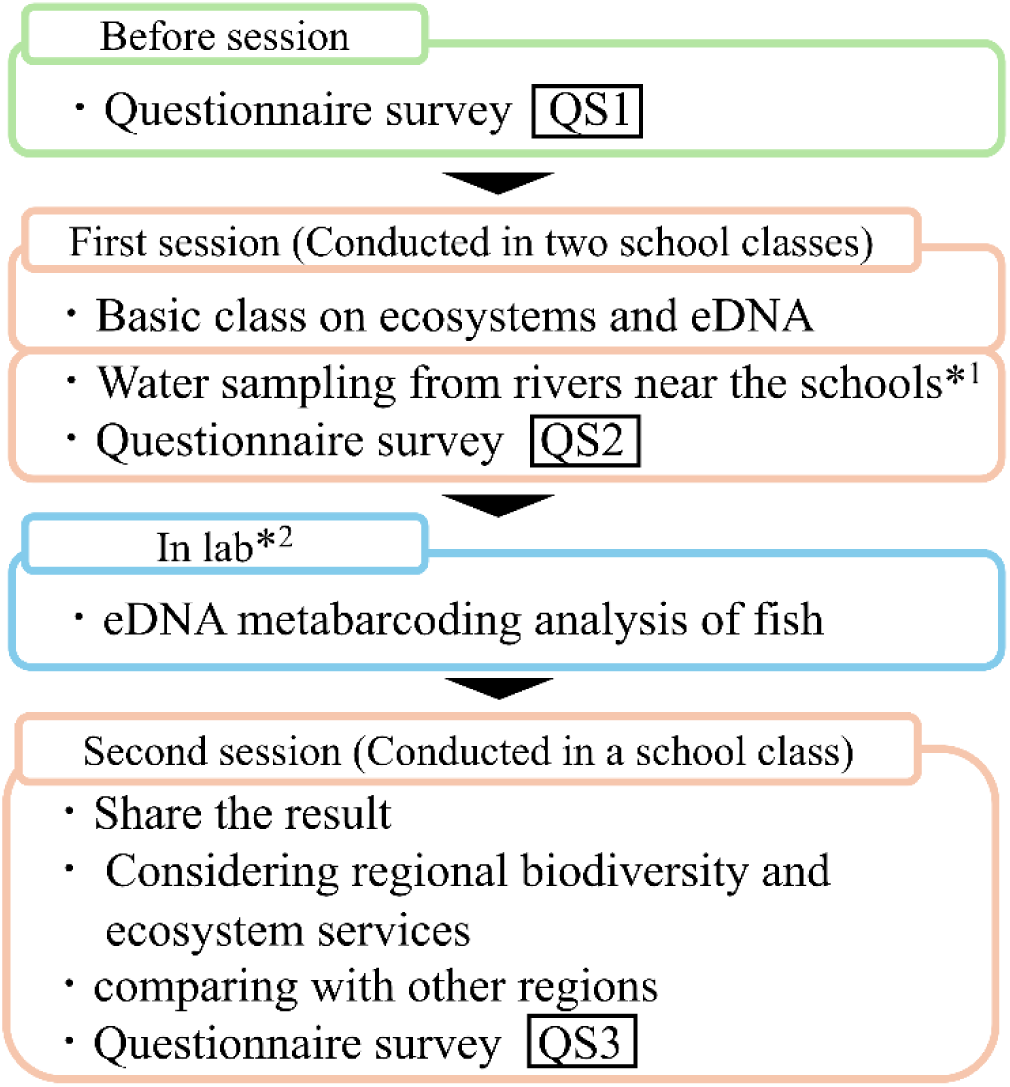
Flowchart of environmental education using eDNA. The lessons were held over 2 d at the school (orange), and each day was scheduled to fit the school’s class time. *1 Second- and third-year students at Yaizuchuo did not participate in this process for reasons related to the school curriculum. This difference was used for comparison while evaluating the effectiveness of the water collection process. * Two students were excluded.

### 2-3. Designing and implementing questionnaires

To evaluate the effectiveness of the program, we conducted three online questionnaire surveys (Tables 2a–c, S1, and S2). The questions were mainly in multiple choice format, with answers provided as either “yes” or “no,” or rating on a five-point Likert scale. We created questions to assess whether the students have intrest about organisms (Table 2a), biodiversity (Table 2b), and ecosystem services (Table 2c). Brief explanations of these ecological terms (Table S1) were provided in QS1 for students without prior knowledge of the concepts. The same questions in QS1, QS2, and QS3 allowed for comparison of the results of the three questionnaires. Questions regarding past nature experiences, based on Sato et al. (2017), were also posed, as differences may exist in educational effectiveness between students with and without rich past nature experiences (Table S2). When students logged into the questionnaire, they were presented with an explanatory statement at the outset. Only those who indicated their agreement were deemed to have provided informed consent and were permitted to proceed. The questionnaire survey was approved by the Ethics Committee of the Graduate School of Human Development and Environment at Kobe University (receipt number: 609-3, Revision-609-3).

**Table 2a.** Questionnaire text regarding likesof organisms.

| No. | Question |
| --- | --- |
| 1 | Do you like biology? |
| 2 | Do you like organisms? |
| 3 | Are you willing to touch organisms? |

**Table 2b.** Questionnaire text regarding biodiversity.

| No. | Question |
| --- | --- |
| 4 | The natural environment (mountains, rivers, sea, satoyama, rice paddies, nature parks, etc.) of my local area is well preserved. |
| 5 | The grounds of the XX High School (the name of the high school where the program was conducted) are rich in nature. |
| 6 | I support prioritizing disaster prevention measures such as riverbank protection work and mountain slope maintenance in my area, even if they negatively impact natural ecosystems. |
| 7 | I support using tax funds to preserve and enrich the diverse natural ecosystems in my local mountains and riparian areas. |
| 8 | The natural environment where I live contains many different species. |
| 9 | Many different species inhabit the river near our high school. |
| 10 | I support removing organisms from my local area if they pose risks to humans. |
| 11 | I support removing organisms from my local area solely because of their strange appearance even if they cause no harm. |
| 12 | I support using tax funds to protect rare species in my local area. |
| 13 | I support using tax funds to remove invasive species in my local area. |
| 14 | I support protecting the local ecosystems by donating money or participating in conservation activities. |
| 15 | I support protecting biodiversity even if it requires sacrificing some modern conveniences (e.g. electricity usage and infrastructure development). |

**Table 2c.** Questionnaire text regarding ecosystem services.

| No. | Question |
| --- | --- |
| 16 | Do you think that the ecosystem services in your local area should be improved? |
| 17 | Do you think that the ecosystem disservices in your local area should be controlled? |
| 18 | Do you think that an increase in biodiversity will also increase ecosystem services? |
| 19 | Do you think that an increase in biodiversity will also increase ecosystem disservices? |

These were used in all the questionnaires: QS1, QS2, and QS3. The survey was conducted in Japanese.

### 2-4. Data processing of survey results

Questionnaire data were divided into two groups: data from students who did collect water, and from those that and did not. Responses to binary questions, such as “yes/no,” were converted to “1/0” data, and responses to questions with a Likert scale were converted to “5/4/3/2/1” from strongest to weakest. To assess the session effects on student attitudes toward biodiversity and ecosystem services/disservices, the analyses included only students who completed all three questionnaires (QS1–QS3). To assess attitudinal changes in students, we coded a rating increase between questionnaires as 1, and no increase as 0; similarly, a rating decrease was coded as 1, and no decrease was coded as 0.

### 2-5. Water sampling by high school students

Students visited rivers near their schools and collected water samples from any chosen location within the designated area of the river. Water sampling details are provided in the Supplementary Material (S1).

### 2-6. eDNA metabarcoding analysis

eDNA was extracted from collected water samples and used for eDNA metabarcoding of fish using MiFish primers (Miya et al., 2015). The obtained data were processed using the MitoFish pipeline (Sato et al., 2018; Zhu et al., 2023). Details of eDNA extraction and bioinformatics are provided in the Supplementary Material (S2‒4).

### 2-7. Data analysis

Statistical analyses were performed using R4.2.1 (R Core Team, 2022). We compared the results of questionnaires QS1, QS2, and QS3 using the Wilcoxon signed‒rank test, which was performed using the wilcox.exact function in the exactRankTests package with Bonferroni correction. We also calculated the effect size *r* using the Z-score and total sample size N, as follows (Mizumoto & Takeuchi, 2008):

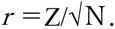

To compare the educational effects between students with and without water sampling experience, we conducted a chi-square test. To test which student characteristics predicted significant increases in interest after the program, we conducted a logistic regression analysis. The binary data for each question processed in Sections 2-2–2-4 served as the dependent variable, and the explanatory variables included questionnaire results regarding students liking of organisms, willingness to touch organisms, and binary data from questions regarding past natural experiences. To reduce the dimensionality of the explanatory variables, we clustered past nature experience data into two groups using the Ward method with squared Euclidean distance (Fig. S1). These clusters were primarily distinguished between students who had few interactions with people knowledgeable about organisms and those who had more interactions (Figs. S1b and c), and we converted these clusters into binary data for use as explanatory variables. Using the same analytical approach, we assessed which student attributes were associated with significant decreases in ratings.

## 3. Results

### 3-1 Implementation of questionnaires

The total number of survey responses was 551, 565, and 541 for QS1, QS2, and QS3, respectively (see Table 1). At the three high schools, the total number of students who completed all the questions in questionnaires QS1, QS2, and QS3 was 426: i.e., 26, 64, and 336 at Hidakamioka, Ena, and Yaizuchuo, respectively (Table 1). Of the 336 students in Yaizuchuo, 200 students collected water samples and 136 students did not.

### 3-2. Result of eDNA metabarcoding analysis

The number of raw reads was 878,461 (the total number of samples was 30, including blanks). These were annotated as 36 taxa (Table S4). In Samples from Hidakamioka, Ena, and Yaizuchuo 9, 16, and 22 fish taxa were detected, respectively. The details of these results are provided in the Supplementary Material (Table S5).

### 3-3 Questionnaire results

#### 3-3-1 Change of preferences for organisms

We examined data from 426 students and found changes in their preferences for organisms. For the question “Do you like biology?,” there was a significant increase in the rating between QS1 and QS2 for students who collected water samples (n = 290) (Wilcoxon signed‒rank test with Bonferroni correction, *p* < 0.05; Table 3). There was a significant increase in rating between QS2 and QS3 among students who did not collect water samples (*p* < 0.05). For the question “Are you willing to touch organisms?,” there was a significant increase in ratings between QS2 and QS3 among students who collected water samples (*p* < 0.05).

**Table 3.** Summary of analysis results for questions regarding preferences for organisms. The water sampling column indicates whether the water was sampled in our program. +indicates that water sampling was performed, and - indicates that water sampling was not performed. QS1 vs. QS2 and QS2 vs. QS3 indicate the questionnaires that were compared. Bold numbers in each cell indicate a significant difference using the Wilcoxon signed‒rank test and Bonferroni correction.

| Question | Water<br>sampling | QS1 vs. QS2 |  | QS2 vs. QS3 |  |
| --- | --- | --- | --- | --- | --- |
| | | Effect size ($r$) | $p$ value | Effect size ($r$) | $p$ value |
| Do you like biology? | + | **0.11** | **< 0.05** | 0.07 | 0.0682 |
|  | - | 0.00 | 1.00 | **0.15** | **< 0.05** |
| Do you like organisms? | + | 0.00 | 1.00 | 0.04 | 0.24 |
|  | - | 0.00 | 1.00 | **0.15** | **< 0.05** |
| Are you willing to<br>touch organisms? | + | 0.00 | 1.00 | **0.11** | **< 0.05** |
|  | - | 0.00 | 1.00 | 0.06 | 0.30 |

Many students showed changes in their response patterns regarding their preferences for organisms (Supplementary Table S6a). However, chi-square tests revealed no significant differences between students with and without water sampling experience for any question.

#### 3-3-2. Change in interest in biodiversity

Student interest in biodiversity changed significantly throughout the program. For the biodiversity questions, we found significant positive changes toward biodiversity in 10 out of 12 questions (Table 4). Between QS1 and QS2, a significant increase in biodiversity interest was observed in seven questions: all changes indicated increased interest in biodiversity (i.e., rating increases in six questions and decreases in one reverse-coded question, both indicating enhanced interest). For the six questions, significant changes were found only for students who collected water samples. “Many different species inhabit the river near our high school” was the only question for which significant rating increases were confirmed for both student with and without water sampling experience. The most significant increase between QS1 and QS2 was found for the questions “I support protecting the local ecosystems by donating money or participating in conservation activities” (Wilcoxon signed‒rank test and Bonferroni correction, *r* = 0.27, *p* < 0.001) and “Many different species inhabit the river near our high school” (*r* = 0.27, *p* < 0.001), followed by “The natural environment where I live contains many different species” (*r* = 0.17, *p* < 0.001). Between QS1 and QS2, the percentages of students whose rating increased between those experienced water sampling and those who did not were significantly different in the three following questions: “I support using tax funds to preserve and enrich the diverse natural ecosystems in my local mountains and riparian areas,” “Many different species inhabit the river near our high school,” and “I support protecting the local ecosystems by donating money or participating in conservation activities” (all these *p*-values were < 0.05: Table S6b).

**Table 4.**
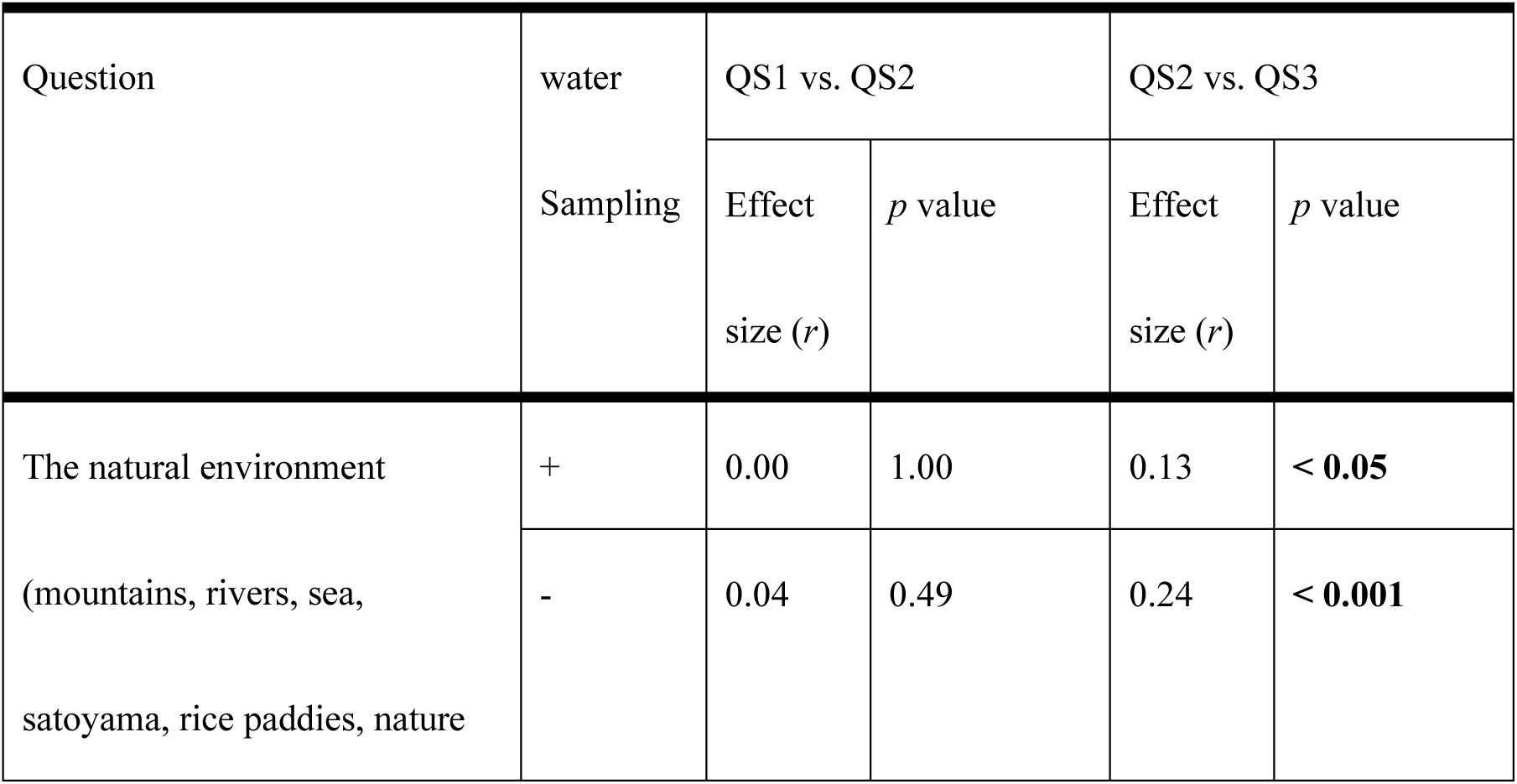
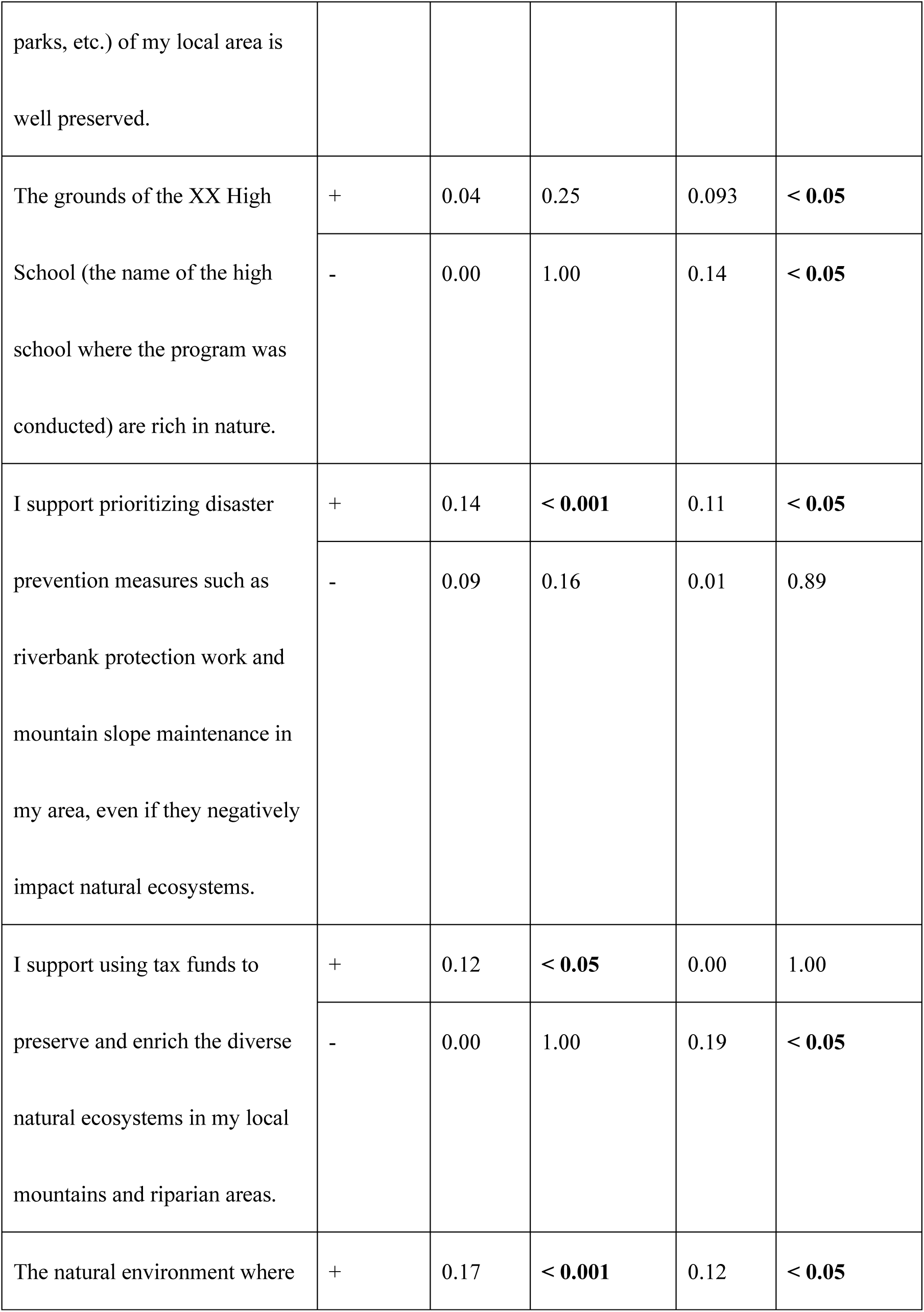
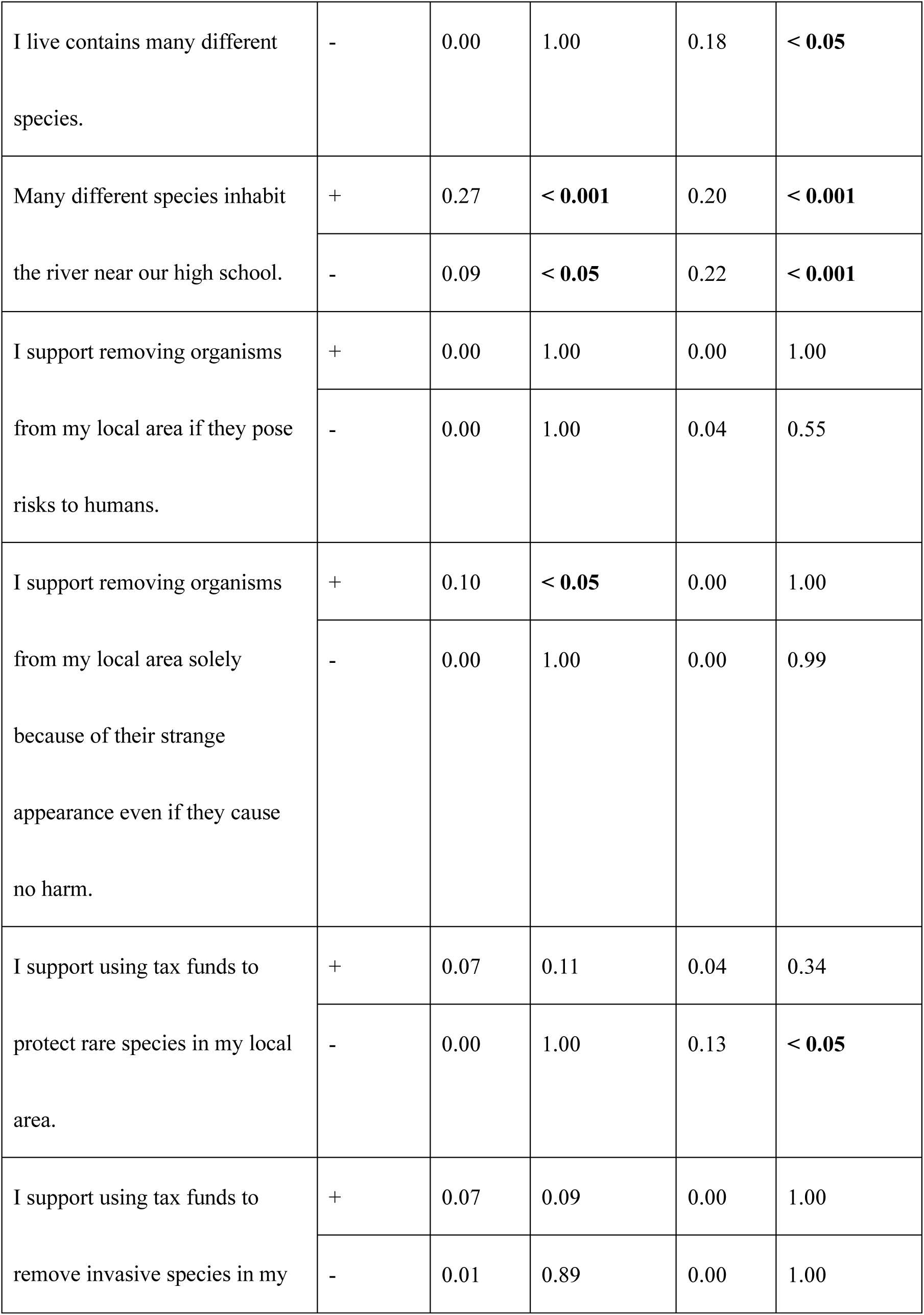
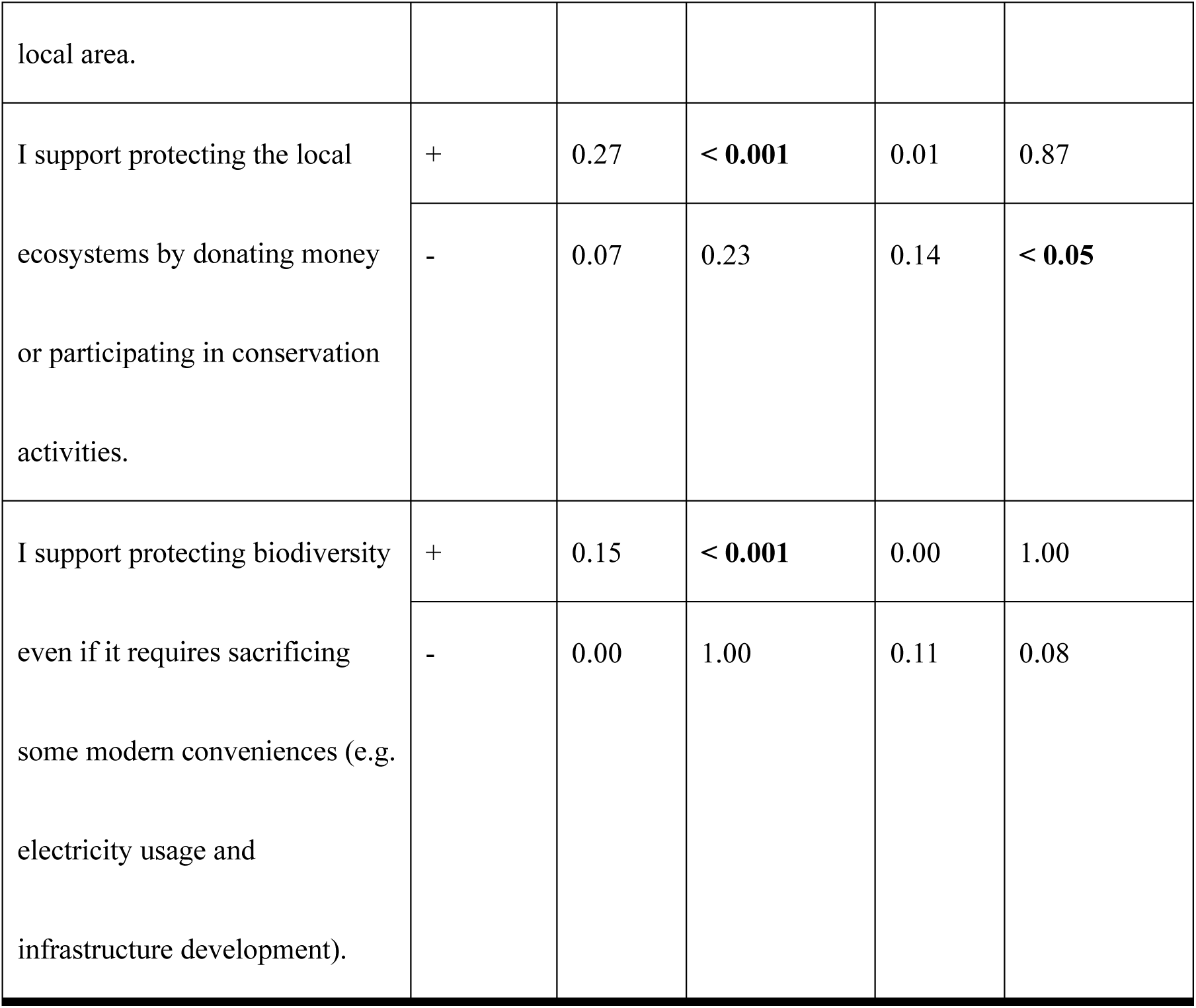
Summary of analysis results for questions regarding biodiversity. The water sampling column indicates whether the water was sampled in our program. + indicates that water sampling was performed, and - indicates that water sampling was not performed. QS1 vs. QS2 and QS2 vs. QS3 indicate the questionnaires that were compared. Bold numbers indicate significant effects found using the Wilcoxon signed‒rank test and Bonferroni correction.

Significant differences were found for eight questions between QS2 and QS3. For most questions, the responses increased the willingness to preserve a high level of biodiversity; however, for one question, the responses changed to prefer a low level of biodiversity. We found significant positive changes toward biodiversity in the three questions for both student groups. For the four questions, significant positive changes were found only for students who did not collect water samples. Furthermore, in one question, “I support prioritizing disaster prevention measures such as riverbank protection work and mountain slope maintenance in my area, even if they negatively impact natural ecosystems,” a significant change to indicate less willingness to preserve biodiversity was found only for the students who collected water. The most significant increase between questions QS2 and QS3 was for students who had experienced water sampling: “Many different species inhabit the river near our high school” (*r* = 0.20, *p* < 0.001). Between QS2 and QS3, we found a significant difference between student groups (*p* < 0.05) in the percentage of students who increased their rating for the question “I support using tax funds to preserve and enrich the diverse natural ecosystems in my local mountains and riparian areas” (Table S6b). Interestingly, the percentage was higher among students who did not collect water than among those who did.

#### 3-3-3 Change in interest in ecosystem services

Through this programme, we found a significant increase in student interest in ecosystem services (Table 5). There was a significant increase in ratings for all four questions between QS1 and QS2 among students with and without water sampling experience. The largest difference was observed for the question “Do you think that the ecosystem services in your local area should be improved?” (Wilcoxon signed‒rank test and Bonferroni correction, *r* = 0.28, *p* < 0.001). Students without water sampling experience exhibited the largest increase in response to the question “Do you think that an increase in biodiversity will increase ecosystem disservices?” (*r* = 0.32, *p* < 0.001). Between QS1 and QS2, we found a significant difference in the percentage of students who increased their rating between students with and without water sampling experience only for the question “Do you think that an increase in biodiversity will also increase ecosystem services?” (chi-square test, *p* < 0.05), such that the percentage was higher for those with water sampling experience than for those without (Table S6c).

**Table 5.** Summary of analysis results for questions regarding ecosystem services and disservices. The water sampling column indicates whether the water was sampled in our program. + indicates that water sampling was performed, and - indicates that water sampling was not performed. QS1 vs. QS2 and QS2 vs. QS3 indicate the questionnaires that were compared. Bold numbers in each cell indicate a significant difference using the Wilcoxon signed‒rank test and Bonferroni correction.

| Question | water | QS1 vs. QS2 |  | QS2 vs. QS3 |  |
| --- | --- | --- | --- | --- | --- |
|  | Sampling | Effect size (*r*) | *p* value | Effect size (*r*) | *p* value |
| Do you think that the ecosystem services in your local area should be improved? | + | 0.28 | < **0.001** | 0.09 | < **0.05** |
|  | - | 0.19 | < **0.05** | 0.12 | < **0.05** |
| Do you think that the ecosystem disservices in your local area should be controlled? | + | 0.25 | < **0.001** | 0.00 | 1.00 |
|  | - | 0.18 | < **0.05** | 0.00 | 1.00 |
| Do you think that an increase in biodiversity will also increase ecosystem services? | + | 0.24 | < **0.001** | 0.02 | 0.66 |
|  | - | 0.18 | < **0.05** | 0.00 | 1.00 |
| Do you think that an increase in biodiversity will also increase ecosystem disservices? | + | 0.22 | < **0.001** | 0.066 | 0.11 |
|  | - | 0.32 | < **0.001** | 0.097 | 0.11 |

Between QS2 and QS3, a significant increase was found only for the question “Do you think that the ecosystem services in your local area should be improved?” for students with and without water sampling experience (*r* = 0.09, *p* < 0.05, and *r* = 0.12, *p* < 0.05, respectively). No significant differences in the percentage of students who increased their ratings were found for any questions between QS2 and QS3.

#### 3-3-4 Student attributes which influenced rating changes following the program

Logistic regression analysis revealed significant effects of student personal attributes on some survey results (Tables 6 and S7). First, we present the results for students who collected water samples. For the questions “I support using tax funds to preserve and enrich the diverse natural ecosystems in my local mountains and riparian areas” and “I support removing organisms from my local area if they pose risks to humans,” students who did not likeorganisms showed a significant increase in their rating (logistic regression analysis; both *p*-values were < 0.05). For the question “I support removing organisms from my local area solely because of their strange appearance even if they cause no harm,” students who liked had a positive opinion of organisms showed a significant increase in their ratings (*p* < 0.05). Students who were willing to touch living organisms significantly developed more balanced perspectives on nature conservation and management for the following questions: “I support using tax funds to preserve and enrich the diverse natural ecosystems in my local mountains and riparian areas,” “The natural environment where I live contains many different species,” and “I support removing organisms from my local area if they pose risks to humans” (all these *p*-values were < 0.05). For the question “I support protecting the local ecosystems by donating money or participating in conservation activities,” students who had few nature experiences showed a significant increase in rating than those who had more s (*p* < 0.05).

Next, we present results for students who did not experience water sampling (Table 6b). For the question “Do you think that an increase in biodiversity will also increase ecosystem services?,” the ratings were significantly lower for students who were unwilling to touch organisms and for those who said they would like to touch organisms (*p* < 0.05). In response to the question “Do you think that an increase in biodiversity will also increase ecosystem disservices?,” students who were not willing to touch organisms significantly increased their ratings, whereas students who were willing to touch organisms and those who had a positive opinion of organisms significantly decreased their ratings (*p* < 0.001, *p* < 0.05, and *p* < 0.001, respectively). For the question “I support protecting the local ecosystems by donating money or participating in conservation activities,” students with more natural experiences exhibited significantly lower ratings (*p* < 0.05).

**Table 6a.**
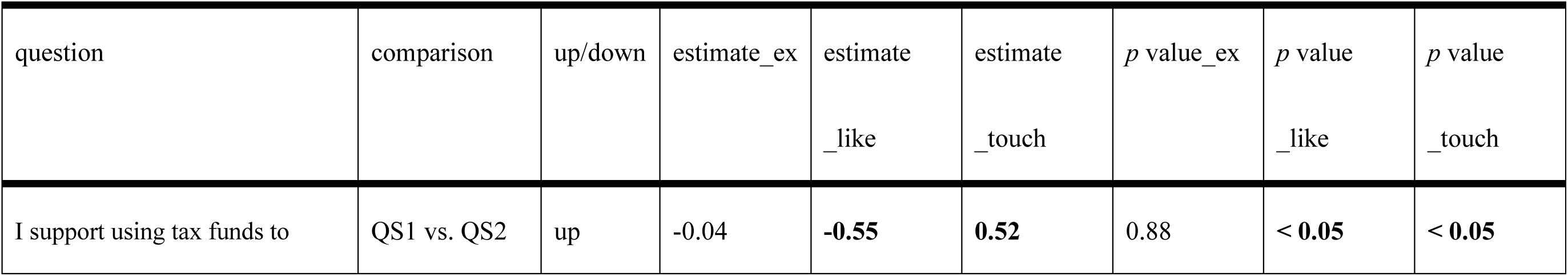
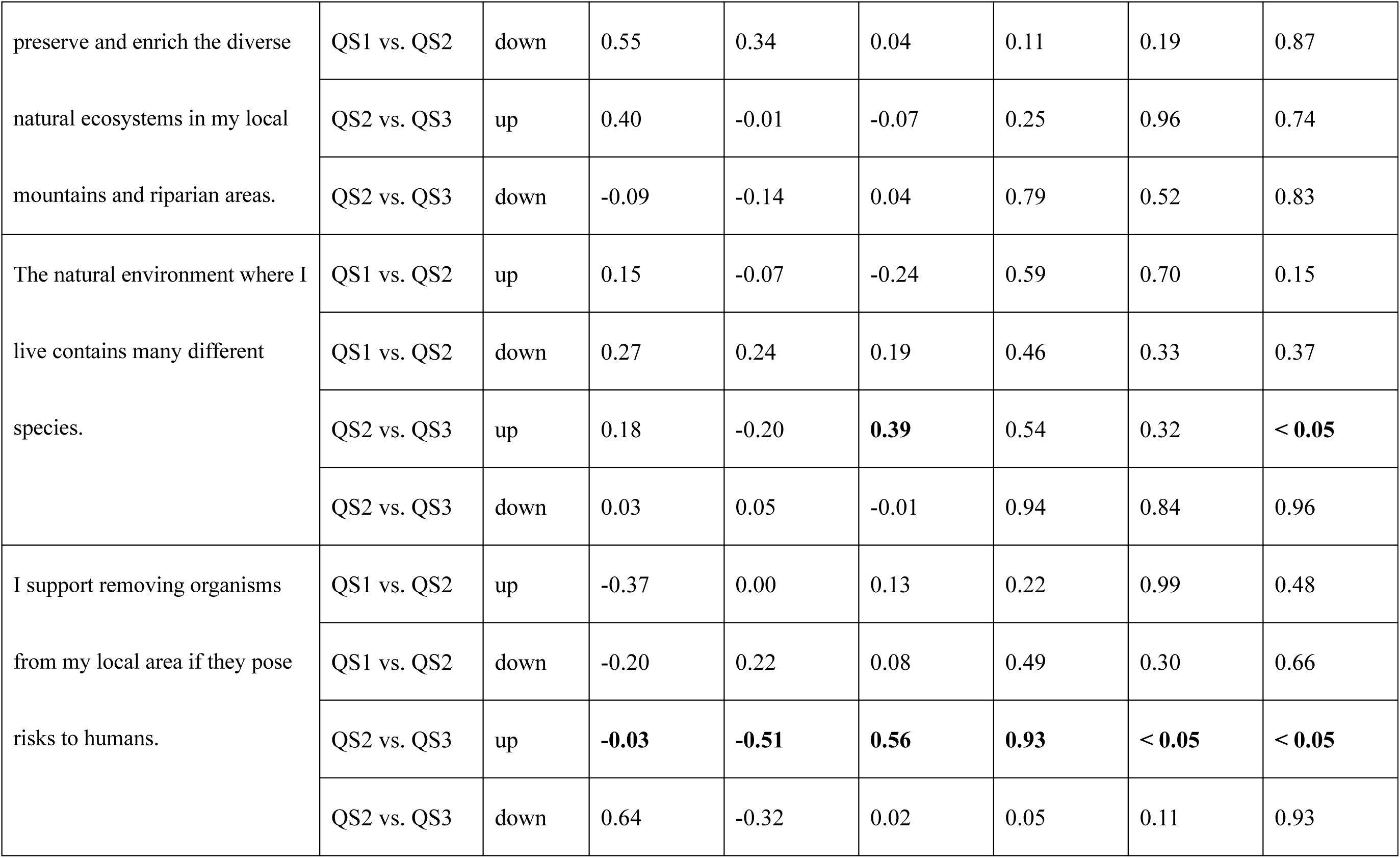
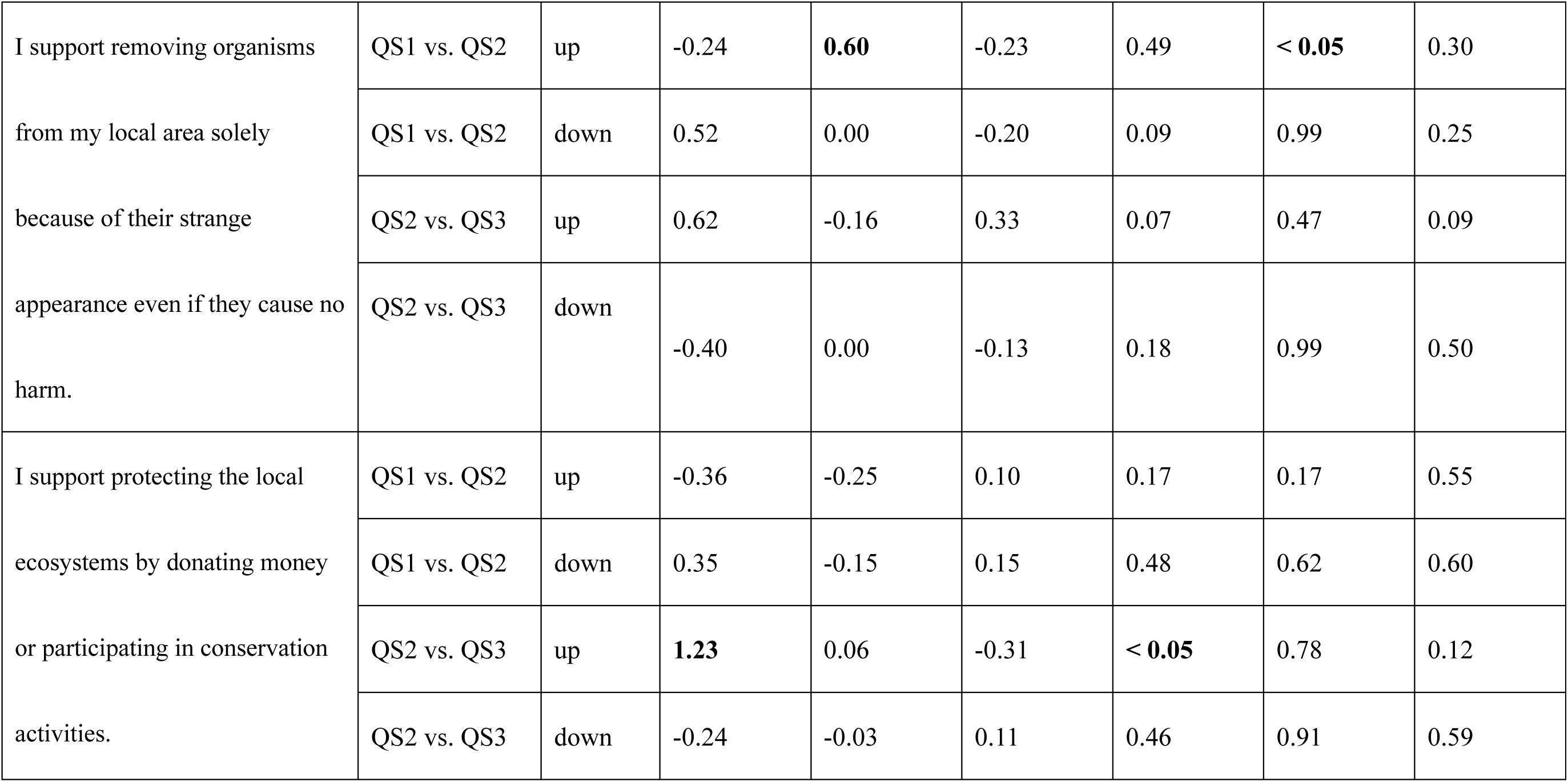
Summary of results of logistic regression analysis for students who experienced water sampling. Each column displays the following information: Question: The text of each question. up/down: Indicates the dependent variable used, where up represents responses that increased between surveys and down represents responses that decreased. Comparison: This indicates that the questionnaires were compared. estimate_ex, like, touch: These show the results of the estimates for the explanatory variables of the student past natural experiences, whether the students had positive opinions of organisms, and whether they wanted to touch them. *p* value_ex, like touch: This shows the significance of the difference in each explanatory variable. Bold text indicates significant differences. Only results for questions with significant differences are listed. For other results, please refer to Table S7.

**Table 6b.**
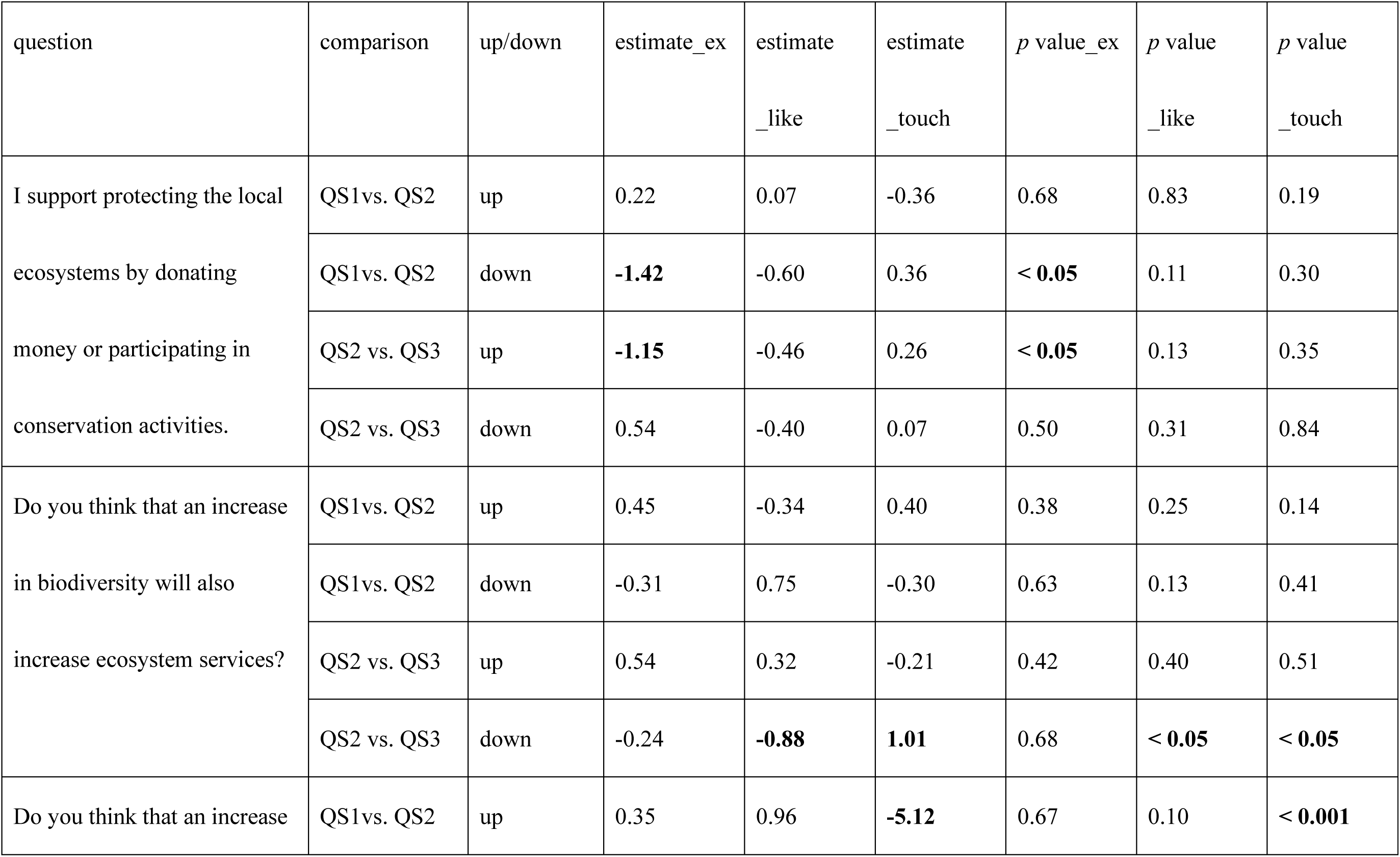
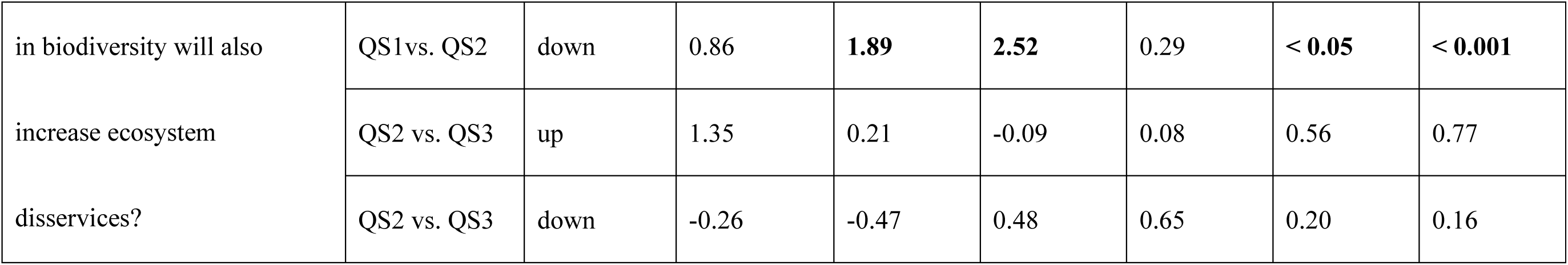
Results of logistic regression analysis of students who did not collect water samples.

| question | comparison | up/down | estimate_ex | estimate<br>_like | estimate<br>_touch | *p* value_ex | *p* value<br>_like | *p* value<br>_touch |
| --- | --- | --- | --- | --- | --- | --- | --- | --- |
| I support protecting the local ecosystems by donating money or participating in conservation activities. | QS1 vs. QS2 | up | 0.22 | 0.07 | -0.36 | 0.68 | 0.83 | 0.19 |
|  | QS1 vs. QS2 | down | **-1.42** | -0.60 | 0.36 | **< 0.05** | 0.11 | 0.30 |
|  | QS2 vs. QS3 | up | **-1.15** | -0.46 | 0.26 | **< 0.05** | 0.13 | 0.35 |
|  | QS2 vs. QS3 | down | 0.54 | -0.40 | 0.07 | 0.50 | 0.31 | 0.84 |
| Do you think that an increase in biodiversity will also increase ecosystem services? | QS1 vs. QS2 | up | 0.45 | -0.34 | 0.40 | 0.38 | 0.25 | 0.14 |
|  | QS1 vs. QS2 | down | -0.31 | 0.75 | -0.30 | 0.63 | 0.13 | 0.41 |
|  | QS2 vs. QS3 | up | 0.54 | 0.32 | -0.21 | 0.42 | 0.40 | 0.51 |
|  | QS2 vs. QS3 | down | -0.24 | **-0.88** | **1.01** | 0.68 | **< 0.05** | **< 0.05** |
| Do you think that an increase | QS1 vs. QS2 | up | 0.35 | 0.96 | **-5.12** | 0.67 | 0.10 | **< 0.001** |
| in biodiversity will also | QS1 vs. QS2 | down | 0.86 | **1.89** | **2.52** | 0.29 | **< 0.05** | **< 0.001** |
| increase ecosystem | QS2 vs. QS3 | up | 1.35 | 0.21 | -0.09 | 0.08 | 0.56 | 0.77 |
| disservices? | QS2 vs. QS3 | down | -0.26 | -0.47 | 0.48 | 0.65 | 0.20 | 0.16 |

## 4. Discussion

### 4-1 Change in preferences for organisms following the eDNA-based educational program

The questionnaire results showed that our eDNA-based educational program improved student attitudes toward biology, nature, and organisms. Specifically, we found that the water sampling experience increased student interest in biology (Table 3). Collecting water samples was the only opportunity in this study for students to interact with the eDNA-based fish survey. Therefore, by being exposed to the simplicity of eDNA analysis, which enables fish diversity identification without direct capture, student interest in biology increased. Water sampling is the only opportunity to directly interact with the natural environment in this program. Direct contact with the natural environment positively affects perceptions of nature (Collado et al., 2013; Soga & Gaston, 2016). Nature experiences through water sampling activities likely increased the positive feelings of students toward local nature and organisms. In addition, students with water sampling experience became more interested in touching organisms throughout the workshop class (*p* < 0.05, water sampling; *p* = 0.30, no water sampling). By confirming the list of local fish species obtained without touching the organisms, student desire to directly touch and observe them increased.

Even for students without water sampling experience, a workshop class improved their interest in biology and organisms, supporting previous research showing the effectiveness of active hands-on learning workshops in biology (Kingdom‒Aaron et al., 2019; Badesaba, 2022).

### 4-2 Changes in interests in biodiversity following the eDNA-based educational program

In 9 of the 12 questions regarding biodiversity, student ratings changed towards affirming the importance of high biodiversity following our program (Table 4). In particular, for the six questions, significant positive changes between QS1 and QS2 were found only among students with water sampling experience. The experience of collecting water for eDNA analysis appeared to have increased student interest in biodiversity; in particular, a significant effect of the experience was observed for the question “I support protecting the local ecosystems by donating money or participating in conservation activities.” According to the Tbilisi Declaration, the objectives of environmental education are to: (1) understand major societal issues; (2) develop skills and attitudes for environmental protection, and; (3) involve people in the problem-solving process and encourage proactive participation. The significant improvements in the above questions showed that our program achieved goal (3). In addition, ratings for the questions “Many different species inhabit the river near our high school” and “The natural environment where I live contains many different species” increased through the program for students who experienced water sampling. The increases indicated that our program helped students understand local biota. Thus, an eDNA-based educational program is a feasible teaching method for high school environmental education.

Conversely, ratings for the question “I support prioritizing disaster prevention measures such as riverbank protection work and mountain slope maintenance in my area, even if they negatively impact natural ecosystems” changed towards accepting potential reductions in biodiversity (Table 4). Since Japan has experienced many natural disasters, such as floods (Naito et al., 2022) and tsunamis (Higaki et al,. 2023), people have developed a culture with a high level of awareness of disaster prevention (Pastrana‒Huguet et al., 2022). Given this national mindset, it can be inferred that viewing the natural landscape during water sampling activities may have strengthened awareness of disaster prevention. Notably, students also showed increased support for “using tax funds to preserve and enrich the diverse natural ecosystems in my local mountains and riparian areas,” indicating that they began to perceive disaster prevention and biodiversity conservation as complementary rather than competing priorities. It is important to emphasise this in the pre-explanation to convey that there are methods for solving the trade-off between ecosystem conservation and disaster prevention, such as sustainable floodplain management for flood prevention (Kiedrzyńska et al., 2015).

Therefore, it is preferable to include a water sampling process when conducting eDNA-based environmental education programs. For several questions, those without water sampling experience showed significant increases in ratings between QS2 and QS3 (Tables 4 and S5b). The workshop itself was shown to enhance student interest in biodiversity and organisms; however, if it were not implemented together with water sampling, the ratings for some questions would have been significantly lower for certain students (Table 6b).

### 4-3 Changes in interests in ecosystem services following the eDNA-based educational program

We found significant changes in student ratings for all four questions related to ecosystem services and disservices through our program (Table 5), indicating its effectiveness in raising awareness regarding ecosystem services and disservices. Rating changes between QS1 and QS2 were found for all questions, regardless of water sampling experience. Additionally, there was a significant increase in the rating of the question “Do you think that the ecosystem services in your local area should be improved?” between QS2 and QS3. We expect that an increased interest in ecosystem services will encourage students to conserve local nature and biodiversity.

Our eDNA-based educational programme also increased student interest in ecosystem services (Table 5). Increased awareness of disservices can prevent people from supporting biodiversity conservation because they are more willing to control disservices than promote services. A balanced understanding of ecosystem services and disservices is essential for informed conservation planning in local communities. Thus, our program appeared to have fostered a balanced perspective among students.

### 4-4 Effectiveness of eDNA-based education for students with low interest in nature

Our results demonstrated that eDNA-based education increased interest in biodiversity even among students with minimal prior interest in biology or nature. Although students who did not have a positive opinion of organisms became more supportive of removing harmful organisms, they also exhibited an increased willingness to use tax funds to protect ecosystems (see Table 6a). Avoiding dangerous organisms is thought to be an innate behaviour that provides a survival advantage (Gullone, 2000), making biophobia difficult to overcome. Furthermore, students with fewer natural experiences showed an increased willingness to donate money or participate in conservation activities following the program (Table 6a). Childhood nature experiences usually facilitate attitudes toward ecosystems and biodiversity conservation (Sato et al., 2017; Aoshima et al., 2023; Barrable et al., 2024). Therefore, our program effectively enhanced the desire of student with low affinities for organisms and nature to protect local ecosystems. If many people have positive opinions of local ecosystems, they can recognise their importance, which will lead to more inclusive conservation with broader citizen participation.

### 4-5 Advantages of eDNA analysis in environmental education

From the river water samples collected by students, 9–22 different fish taxa were identified (Tables S5a‒c). In principle, the process of catching and identifying many fish species requires substantial time and labour expenses. In particular, when this is conducted as part of environmental education, these costs are likely to be higher because of the lack of specialised experts and limited time. These limitations are serious in educational situations during class. Controversially, eDNA analysis has the advantage of identifying a range of species at a lower cost than conventional survey methods (Kelly et al., 2014). We successfully collected water samples by travelling from the school to the river and back in < 1 h. The strength of eDNA analysis, with its low time and labour costs, is also considered a major advantage when used in environmental education of biodiversity.

Another advantage of eDNA analysis is that it can be performed in a variety of environments. In environmental education, the challenge is that the organisms that can be used as research subjects depend on their locations (Iwanishi & Takada, 2016). Because eDNA analysis is not limited to specific target species, results that reflect local biota can be obtained. Additionally, if there are no water bodies in the implementation area, eDNA sampling can be conducted from sources other than aquatic environments. For example, previous studies have reported soil and air eDNA analyses of taxa other than fish (Leempoel et al., 2020; Warmer et al., 2024). Therefore, it is possible to conduct environmental education using eDNA analysis in a variety of areas by selecting the target taxa and sampling methods for study sites. However, it is unclear whether the same educational effect can be achieved by changing the methods; thus, further verification is required.

## 5. Conclusion

Our eDNA-based education program in high school classes successfully improved student awareness of nature and organisms and increased their interest in biodiversity and ecosystem services. In addition, multiple questionnaire surveys revealed that the experience of water sampling for eDNA analysis combined with workshop activities using the analysis results effectively enhanced student connections to local biodiversity. The results of this study obtained from a wide range of high school students will be valuable for future applications of eDNA-based educational programs in environmental education. Because only a limited number of studies have examined the effectiveness of eDNA analyses in citizen education, this field remains in its infancy. We hope that eDNA-based environmental education approaches will expand beyond high schools to include diverse educational settings and communities, facilitating broader engagement with biodiversity conservation across various societies.

## Supporting information

SuppInfo_Fig and Table

SuppInfo_TableS4

## AUTHOR CONTRIBUTIONS

RPK conceived of the study and wrote the first draft of the manuscript. RPK drafted the questionnaire and MKi, MS, AU, and TM improved it by providing critical feedback. RPK, TS, and MKa conducted the program and completed the questionnaires at high schools. RPK conducted eDNA analysis and interpreted the results using TM. RPK, TS, and MKa discussed the method of analysing the questionnaire results and performed data analysis. All the authors provided critical feedback on the manuscript and approved the final version for publication.

## ACKNOWLEDGEMENTS

We thank Mr. Yuto Tanaka of Hidakamioka High School, Mr. Taiki Niwa of Ena High School, and Dr. Yuichi Yaoi of Yaizuchuo High School for supporting the high school environmental education program. We would also like to thank all high school students who participated in the environmental education and questionnaire surveys conducted as part of this research. This study was supported by JSPS KAKENHI (JP22H03813, 23K25067, and 19J00864), JST SPRING (JPMJSP2148), the Kurita Water and Environment Foundation and the Research Grant from the Foundation of Environmental Education Japan(23H024), and the Kobe University 2023 Student Community Action Plan. We used the AI tools DeepL and DeepLwrite to translate and review parts of this study in English. We would like to thank Editage (www.editage.jp) for English language editing.

## Data availability statement

This paper has not yet been published in the archive.

