## Supplementary material for "Feasibility of an eDNA-based educational program for high school students": SuppInfo_Fig and Table

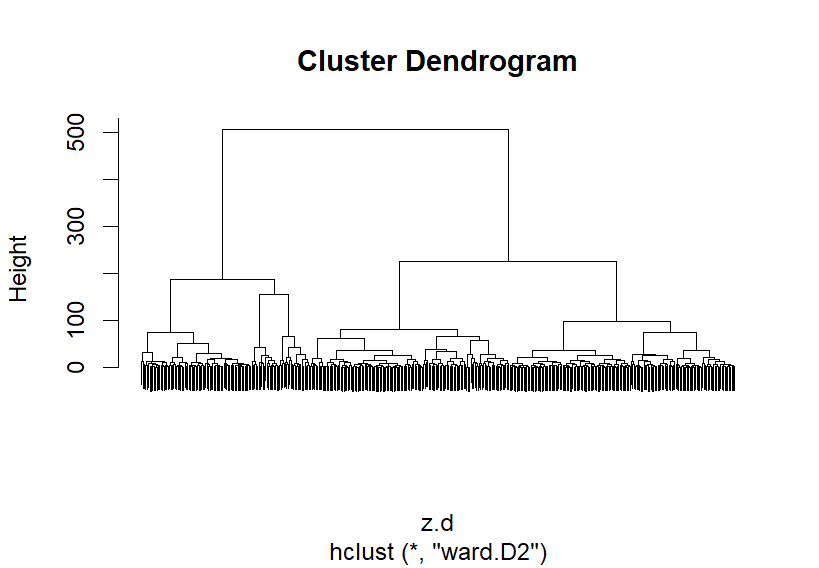

Cluster 1

Cluster 2

**(a)**

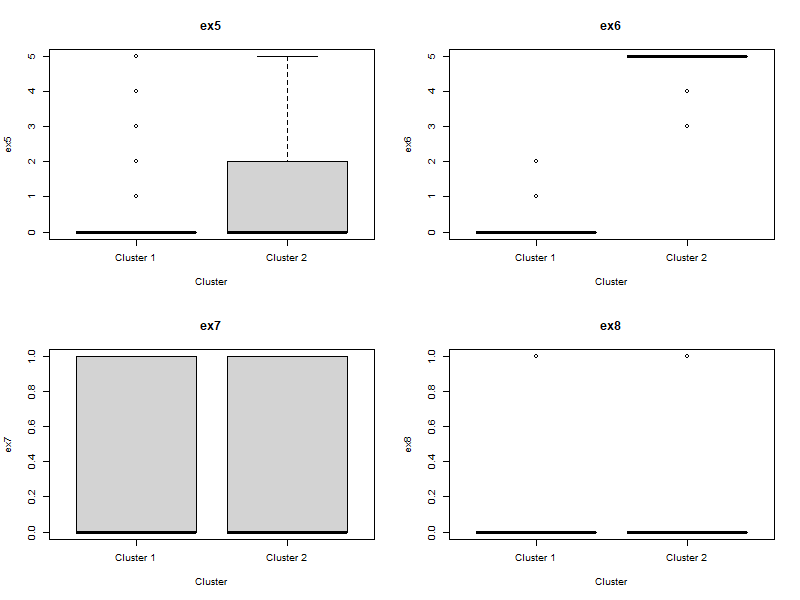

**(b)**

**(c)**

Fig. S1　Cluster analysis of results of questionnaire regarding past nature experiences. (a) Students were clustered using the Ward method based on their responses to questions regarding their past natural experiences. (b) and (c) Questionnaire results for each cluster. Question (b) was: “Did you interact with people who were knowledgeable about nature, such as museum staff or people who were knowledgeable about organisms, before you were in junior high school?” and question (c) was “Did you interact with people who were knowledgeable about nature, such as museum staff or people who were knowledgeable about organisms, in the last few years?.”

Table S1 Questionnaire texts regarding knowledge.

| Instructions | The following questions relate to ecosystems. After reading the these explanations, please select the option that best applies to each question.  ・Biodiversity is the total of genes, species, and ecosystems in a particular area. It refers to the diverse existence of organisms and the places (ecosystems) where they live.  ・Ecosystem services are the benefits that organisms and ecosystems provide to us. These include the supply of food and resources, the absorption of carbon dioxide through photosynthesis, mountain climbing and swimming, nutrient cycling and soil formation, etc.  ・Ecosystem disservices are the opposite of ecosystem services, and are the negative impacts that organisms and ecosystems have on us. These include damage to crops caused by wild animals, damage caused by toxic organisms, infectious diseases, hay fever, etc. |
| --- | --- |
| No. | Question |
| s1 | Did you know what the term “biodiversity” means? |
| s2 | Did you know what the term “ecosystem services” means? |
| s3 | Did you know what the term “ecosystem disservices” means? |

Table S2 Questionnaire texts regarding past natural experiences.

| No. | Question |
| --- | --- |
| s5 | Before entering junior high school, did you have experience hiking mountain areas (mountain trails, forests, mountain streams, nature parks, etc.) or spending time in the waterside environments (the sea, rivers, wetlands, etc.)? |
| s6 | This question is for those who answered yes to s5. How often did you visit mountain or the waterside areas? |
| s7 | In the last couple of years, did you visit the mountains (mountain trails, forests, mountain streams, nature parks, etc.) or to places near the waterside (the sea, rivers, wetlands, etc.)? |
| s8 | This question is for those who answered yes to s7. How often did you visit mountain or waterside areas? |
| s9 | Before entering junior high school, did you take care of animals or insects, or grow plants? |
| s10 | In the last couple of years, have you taken care of animals or insects, or grown plants? |
| s11 | Before entering junior high school, did you interact with people who were nature specialists, such as museum staff or adults who were knowledgeable about organisms? |
| s12 | This is a question for those who answered “yes” to s11. How often did you interact with nature specialist? |
| s13 | In the last couple of years, have you interacted with people who were nature specialists, such as museum staff or adults who were knowledgeable about organisms? |
| s14 | This is a question for those who answered “yes” to s13. How often have you interacted with nature specialists? |
| s15 | Before entering junior high school, did you experience any negative effects from nature　 (ecosystem disservice), such as being attacked by wild animals or insects, or having your garden or flowerbeds damaged? |
| s16 | In the last couple of years, have you experienced any negative effects from nature (ecosystem disservice)? |

Note that these questions were provided to students in Japanese, and answers were provided in a choice format.

Table S3a. Results of questionnaire regarding preferences for biology in all high schools.

| Question | number of questionnaires | water Sampling | total number of responses. |
| --- | --- | --- | --- |
| Do you like biology? | QS1 | + | Ⅰ:1, Ⅱ:29, Ⅲ:115, Ⅳ:112, Ⅴ:33 |
|  |  | - | Ⅰ:2, Ⅱ:19, Ⅲ:36, Ⅳ:61, Ⅴ:18 |
|  | QS2 | + | Ⅰ:3, Ⅱ:23, Ⅲ:89, Ⅳ:139, Ⅴ:36 |
|  |  | - | Ⅰ:2, Ⅱ:20, Ⅲ:28, Ⅳ:69, Ⅴ:17 |
|  | QS3 | + | Ⅰ:1, Ⅱ:22, Ⅲ:81, Ⅳ:141, Ⅴ:45 |
|  |  | - | Ⅰ:2, Ⅱ:14, Ⅲ:29, Ⅳ:63, Ⅴ:28 |
| Do you like organisms? | QS1 | + | Ⅰ:9, Ⅱ:25, Ⅲ:72, Ⅳ:121, Ⅴ:63 |
|  |  | - | Ⅰ:2, Ⅱ:14, Ⅲ:21, Ⅳ:63, Ⅴ:36 |
|  | QS2 | + | Ⅰ:3, Ⅱ:28, Ⅲ:85, Ⅳ:112, Ⅴ:62 |
|  |  | - | Ⅰ:1, Ⅱ:15, Ⅲ:29, Ⅳ:55, Ⅴ:36 |
|  | QS3 | + | Ⅰ:5, Ⅱ:27, Ⅲ:70, Ⅳ:116, Ⅴ:72 |
|  |  | - | Ⅰ:2, Ⅱ:10, Ⅲ:19, Ⅳ:63, Ⅴ:42 |
| Are you willing to touch organisms? | QS1 | + | Ⅰ:26, Ⅱ:52, Ⅲ:102, Ⅳ:81, Ⅴ:29 |
|  |  | - | Ⅰ:8, Ⅱ:20, Ⅲ:37, Ⅳ:48, Ⅴ:23 |
|  | QS2 | + | Ⅰ:14, Ⅱ:69, Ⅲ:101, Ⅳ:78, Ⅴ:28 |
|  |  | - | Ⅰ:7, Ⅱ:22, Ⅲ:37, Ⅳ:43, Ⅴ:27 |
|  | QS3 | + | Ⅰ:13, Ⅱ:54, Ⅲ:99, Ⅳ:92, Ⅴ:32 |
|  |  | - | Ⅰ:3, Ⅱ:21, Ⅲ:38, Ⅳ:47, Ⅴ:27 |

Table S3b. Results of questionnaire regarding preferences for biodiversity in all high schools.

| Question | number of questionnaires | water Sampling | total number of responses. |
| --- | --- | --- | --- |
| The natural environment (mountains, rivers, sea, satoyama, rice paddies, nature parks, etc.) of my local area is well preserved. | QS1 | + | Ⅰ:2, Ⅱ:29, Ⅲ:87, Ⅳ143, Ⅴ:29 |
|  |  | - | Ⅰ:2, Ⅱ:28, Ⅲ:41, Ⅳ:56, Ⅴ:9 |
|  | QS2 | + | Ⅰ:0, Ⅱ:21, Ⅲ:91, Ⅳ:164, Ⅴ:14 |
|  |  | - | Ⅰ:0, Ⅱ:21, Ⅲ:43, Ⅳ:64, Ⅴ:8 |
|  | QS3 | + | Ⅰ:0, Ⅱ:13, Ⅲ:83, Ⅳ:156, Ⅴ:38 |
|  |  | - | Ⅰ:0, Ⅱ:7, Ⅲ:37, Ⅳ:77, Ⅴ:15 |
| The grounds of the XX High School (the name of the high school where the program was conducted) are rich in nature. | QS1 | + | Ⅰ:6, Ⅱ:45, Ⅲ:113, Ⅳ:106, Ⅴ:20 |
|  |  | - | Ⅰ:1, Ⅱ:29, Ⅲ:60, Ⅳ:39, Ⅴ:7 |
|  | QS2 | + | Ⅰ:4, Ⅱ:50, Ⅲ:91, Ⅳ:121, Ⅴ:24 |
|  |  | - | Ⅰ:4, Ⅱ:34, Ⅲ:47, Ⅳ:47, Ⅴ:4 |
|  | QS3 | + | Ⅰ:1, Ⅱ:45, Ⅲ:84, Ⅳ:130, Ⅴ:30 |
|  |  | - | Ⅰ:1, Ⅱ:33, Ⅲ:44, Ⅳ:45, Ⅴ:11 |
| I support prioritizing disaster prevention measures such as riverbank protection work and mountain slope maintenance in my area, even if they negatively impact natural ecosystems. | QS1 | + | Ⅰ:16, Ⅱ:96, Ⅲ:128, Ⅳ:37, Ⅴ:13 |
|  |  | - | Ⅰ:6, Ⅱ:36, Ⅲ:57, Ⅳ:33, Ⅴ:4 |
|  | QS2 | + | Ⅰ:8, Ⅱ:80, Ⅲ:124, Ⅳ:61, Ⅴ:17 |
|  |  | - | Ⅰ:4, Ⅱ:34, Ⅲ:46, Ⅳ:45, Ⅴ:7 |
|  | QS3 | + | Ⅰ:5, Ⅱ:63, Ⅲ:117, Ⅳ:84, Ⅴ:21 |
|  |  | - | Ⅰ:3, Ⅱ:33, Ⅲ:44, Ⅳ:45, Ⅴ:11 |
| I support using tax funds to preserve and enrich the diverse natural ecosystems in my local mountains and riparian areas. | QS1 | + | Ⅰ:3, Ⅱ:24, Ⅲ:125, Ⅳ:112, Ⅴ:26 |
|  |  | - | Ⅰ:0, Ⅱ:8, Ⅲ:46, Ⅳ:65, Ⅴ:17 |
|  | QS2 | + | Ⅰ:4, Ⅱ:14, Ⅲ:94, Ⅳ:155, Ⅴ:23 |
|  |  | - | Ⅰ:0, Ⅱ:7, Ⅲ:46, Ⅳ:67, Ⅴ:16 |
|  | QS3 | + | Ⅰ:2, Ⅱ:25, Ⅲ:86, Ⅳ150, Ⅴ:27 |
|  |  | - | Ⅰ:0, Ⅱ:6, Ⅲ:28, Ⅳ:80, Ⅴ:22 |
| The natural environment where I live contains many different species. | QS1 | + | Ⅰ:3, Ⅱ:42, Ⅲ:87, Ⅳ:118, Ⅴ:40 |
|  |  | - | Ⅰ:0, Ⅱ:27, Ⅲ:23, Ⅳ:69, Ⅴ:17 |
|  | QS2 | + | Ⅰ:0, Ⅱ:19, Ⅲ:66, Ⅳ:174, Ⅴ:31 |
|  |  | - | Ⅰ:0, Ⅱ:14, Ⅲ:33, Ⅳ:81, Ⅴ:8 |
|  | QS3 | + | Ⅰ:1, Ⅱ:10, Ⅲ:56, Ⅳ:175, Ⅴ:48 |
|  |  | - | Ⅰ:0, Ⅱ:9, Ⅲ:24, Ⅳ:82, Ⅴ:21 |
| Many different species inhabit the river near our high school. | QS1 | + | Ⅰ:3, Ⅱ:20, Ⅲ:163, Ⅳ:86, Ⅴ:18 |
|  |  | - | Ⅰ:0, Ⅱ:23, Ⅲ:63, Ⅳ:42, Ⅴ:8 |
|  | QS2 | + | Ⅰ:0, Ⅱ:21, Ⅲ:81, Ⅳ:156, Ⅴ:32 |
|  |  | - | Ⅰ:0, Ⅱ:14, Ⅲ:49, Ⅳ:66, Ⅴ:7 |
|  | QS3 | + | Ⅰ:0, Ⅱ:13, Ⅲ:39, Ⅳ:190, Ⅴ:48 |
|  |  | - | Ⅰ:0, Ⅱ:10, Ⅲ:21, Ⅳ:86, Ⅴ:19 |
| I support removing organisms from my local area if they pose risks to humans. | QS1 | + | Ⅰ:6, Ⅱ:24, Ⅲ:98, Ⅳ:94, Ⅴ:68 |
|  |  | - | Ⅰ:1, Ⅱ:14, Ⅲ:33, Ⅳ:52, Ⅴ:36 |
|  | QS2 | + | Ⅰ:1, Ⅱ:23, Ⅲ:105, Ⅳ:109, Ⅴ:52 |
|  |  | - | Ⅰ:1, Ⅱ:16, Ⅲ:32, Ⅳ:53, Ⅴ:34 |
|  | QS3 | + | Ⅰ:1, Ⅱ:28, Ⅲ:103, Ⅳ:107, Ⅴ:51 |
|  |  | - | Ⅰ:2, Ⅱ:16, Ⅲ:33, Ⅳ:60, Ⅴ:25 |
| I support removing organisms from my local area solely because of their strange appearance even if they cause no harm. | QS1 | + | Ⅰ:15, Ⅱ:67, Ⅲ:94, Ⅳ:75, Ⅴ:39 |
|  |  | - | Ⅰ:8, Ⅱ:36, Ⅲ:39, Ⅳ:31, Ⅴ:22 |
|  | QS2 | + | Ⅰ:18, Ⅱ:68, Ⅲ:111, Ⅳ:72, Ⅴ:21 |
|  |  | - | Ⅰ:6, Ⅱ:41, Ⅲ:33, Ⅳ:40, Ⅴ:16 |
|  | QS3 | + | Ⅰ:15, Ⅱ:75, Ⅲ:111, Ⅳ:61, Ⅴ:28 |
|  |  | - | Ⅰ:8, Ⅱ:41, Ⅲ:34, Ⅳ:39, Ⅴ:14 |
| I support using tax funds to protect rare species in my local area. | QS1 | + | Ⅰ:2, Ⅱ:19, Ⅲ:102, Ⅳ:129, Ⅴ:38 |
|  |  | - | Ⅰ:0, Ⅱ:9, Ⅲ:34, Ⅳ:71, Ⅴ:22 |
|  | QS2 | + | Ⅰ:4, Ⅱ:12, Ⅲ:89, Ⅳ:140, Ⅴ:45 |
|  |  | - | Ⅰ:0, Ⅱ:11, Ⅲ:32, Ⅳ:72, Ⅴ:21 |
|  | QS3 | + | Ⅰ:2, Ⅱ:9, Ⅲ:88, Ⅳ:138, Ⅴ:52 |
|  |  | - | Ⅰ:0, Ⅱ:6, Ⅲ:28, Ⅳ:72, Ⅴ:30 |
| I support using tax funds to remove invasive species in my local area. | QS1 | + | Ⅰ:2, Ⅱ:24, Ⅲ:122, Ⅳ:116, Ⅴ:26 |
|  |  | - | Ⅰ:0, Ⅱ:11, Ⅲ:39, Ⅳ:69, Ⅴ:17 |
|  | QS2 | + | Ⅰ:1, Ⅱ:24, Ⅲ:102, Ⅳ:133, Ⅴ:30 |
|  |  | - | Ⅰ:0, Ⅱ:6, Ⅲ:42, Ⅳ:67, Ⅴ:21 |
|  | QS3 | + | Ⅰ:3, Ⅱ:21, Ⅲ:99, Ⅳ:135, Ⅴ:32 |
|  |  | - | Ⅰ:0, Ⅱ:6, Ⅲ:36, Ⅳ:72, Ⅴ:30 |
| I support protecting the local ecosystems by donating money or participating in conservation activities. | QS1 | + | Ⅰ:12, Ⅱ:55, Ⅲ:134, Ⅳ:80, Ⅴ:9 |
|  |  | - | Ⅰ:0, Ⅱ:19, Ⅲ:48, Ⅳ:62, Ⅴ:7 |
|  | QS2 | + | Ⅰ:4, Ⅱ:38, Ⅲ:112, Ⅳ:122, Ⅴ:14 |
|  |  | - | Ⅰ:0, Ⅱ:16, Ⅲ:39, Ⅳ:74, Ⅴ:7 |
|  | QS3 | + | Ⅰ:6, Ⅱ:28, Ⅲ:113, Ⅳ:132, Ⅴ:11 |
|  |  | - | Ⅰ:0, Ⅱ:8, Ⅲ:36, Ⅳ:83, Ⅴ:9 |
| I support protecting biodiversity even if it requires sacrificing some modern conveniences (e.g. electricity usage and infrastructure development). | QS1 | + | Ⅰ:13, Ⅱ:74, Ⅲ:153, Ⅳ:42, Ⅴ:8 |
|  |  | - | Ⅰ:2, Ⅱ:41, Ⅲ:61, Ⅳ:28, Ⅴ:4 |
|  | QS2 | + | Ⅰ:6, Ⅱ:57, Ⅲ:149, Ⅳ:67, Ⅴ:11 |
|  |  | - | Ⅰ:1, Ⅱ:43, Ⅲ:54, Ⅳ:33, Ⅴ:5 |
|  | QS3 | + | Ⅰ:6, Ⅱ:52, Ⅲ:149, Ⅳ:71, Ⅴ:12 |
|  |  | - | Ⅰ:2, Ⅱ:33, Ⅲ:47, Ⅳ:45, Ⅴ:9 |

Table S3c. Results of questionnaire regarding preferences for ecosystem service in all high schools.

| Question | number of questionnaires | water Sampling | total number of responses. |
| --- | --- | --- | --- |
| Do you think that the ecosystem services in your local area should be improved? | QS1 | + | Ⅰ:1, Ⅱ:18, Ⅲ:143, Ⅳ:112, Ⅴ:16 |
|  |  | - | Ⅰ:0, Ⅱ:1, Ⅲ:55, Ⅳ:69, Ⅴ:11 |
|  | QS2 | + | Ⅰ:1, Ⅱ:6, Ⅲ:96, Ⅳ:150, Ⅴ:37 |
|  |  | - | Ⅰ:0, Ⅱ:2, Ⅲ:35, Ⅳ:80, Ⅴ:19 |
|  | QS3 | + | Ⅰ:0, Ⅱ:8, Ⅲ:73, Ⅳ:160, Ⅴ:49 |
|  |  | - | Ⅰ:0, Ⅱ:3, Ⅲ:23, Ⅳ:81, Ⅴ:29 |
| Do you think that the ecosystem disservices in your local areashould be controlled? | QS1 | + | Ⅰ:1, Ⅱ:18, Ⅲ:173, Ⅳ:86, Ⅴ:12 |
|  |  | - | Ⅰ:1, Ⅱ:6, Ⅲ:59, Ⅳ:65, Ⅴ:5 |
|  | QS2 | + | Ⅰ:1, Ⅱ:20, Ⅲ:152, Ⅳ:98, Ⅴ:129 |
|  |  | - | Ⅰ:1, Ⅱ:6, Ⅲ:58, Ⅳ:65, Ⅴ:6 |
|  | QS3 | + | Ⅰ:1, Ⅱ:21, Ⅲ:103, Ⅳ:129, Ⅴ:36 |
|  |  | - | Ⅰ:0, Ⅱ:5, Ⅲ:39, Ⅳ:71Ⅴ:21 |
| Do you think that an increase in biodiversity will also increase ecosystem services? | QS1 | + | Ⅰ:0, Ⅱ:8, Ⅲ:66, Ⅳ:185, Ⅴ:31 |
|  |  | - | Ⅰ:0, Ⅱ:1, Ⅲ:23, Ⅳ:90, Ⅴ:22 |
|  | QS2 | + | Ⅰ:0, Ⅱ:4, Ⅲ:33, Ⅳ:189, Ⅴ:64 |
|  |  | - | Ⅰ:0, Ⅱ:0, Ⅲ:12, Ⅳ:89, Ⅴ:35 |
|  | QS3 | + | Ⅰ:0, Ⅱ:3, Ⅲ:36, Ⅳ:171, Ⅴ:80 |
|  |  | - | Ⅰ:0, Ⅱ:1, Ⅲ:10, Ⅳ:89, Ⅴ:36 |
| Do you think that an increase in biodiversity will also increase ecosystem disservices? | QS1 | + | Ⅰ:0, Ⅱ:7, Ⅲ:86, Ⅳ:166, Ⅴ:31 |
|  |  | - | Ⅰ:0, Ⅱ:6, Ⅲ:34, Ⅳ:78, Ⅴ:18 |
|  | QS2 | + | Ⅰ:0, Ⅱ:3, Ⅲ:44, Ⅳ:194, Ⅴ:49 |
|  |  | - | Ⅰ:0, Ⅱ:2, Ⅲ:12, Ⅳ:97, Ⅴ:25 |
|  | QS3 | + | Ⅰ:0, Ⅱ:0, Ⅲ:43, Ⅳ:183, SⅤ:64 |
|  |  | - | Ⅰ:0, Ⅱ:1, Ⅲ:11, Ⅳ:99Ⅴ:25 |

The water sampling column indicates whether the water was sampled in our program. + indicates that water sampling was performed, and - indicates that water sampling was not performed. The numbers I, II, III, IV, and V in the total number of responses column represent the response options: “strongly disagree.” “disagree,” “neither agree nor disagree,” “agree,” and “strongly agree,” respectively.

Table S5a. Results of eDNA metabarcoding from the Hidakamioka sample.

| Scientific name | reads (1) | reads (2) |
| --- | --- | --- |
| *Phoxinus oxycephalus jouyi* | 16048 | 6877 |
| *Rhinogobius flumineus* | 9869 | 4536 |
| *Pseudaspius hakonensis* | 10241 | 12160 |
| *Oncorhynchus masou masou* | 5544 | 9641 |
| *Plecoglossus altivelis altivelis* | 546 | 1283 |
| *Salvelinus leucomaenis imbrius* | 0 | 2610 |
| *Cottus pollux* | 0 | 2220 |
| *Salvelinus leucomaenis pluvius* | 0 | 1388 |
| *Niwaella delicata* | 0 | 677 |
| *Homo sapiens* | 275 | 176 |

Table S5b. Results of eDNA metabarcoding from the Ena sample.

| Scientific name | reads (1) | reads (2) | reads (3) |
| --- | --- | --- | --- |
| *Phoxinus steindachneri* | 13517 | 6235 | 1642 |
| *Rhinogobius flumineus* | 8975 | 5790 | 4758 |
| *Rhinogobius sp.* | 3059 |  | 3418 |
| *Phoxinus oxycephalus jouyi* | 2240 | 4620 | 1069 |
| *Cyprinus carpio* | 4684 | 2801 | 397 |
| *Zacco platypus* | 2280 | 2244 | 11354 |
| *Liobagrus reinii* | 1970 | 801 |  |
| *Plecoglossus altivelis altivelis* | 1661 |  |  |
| *Opsariichthys uncirostris uncirostris* | 1697 |  |  |
| *Carassius sp.* |  | 520 |  |
| *Carassius cuvieri* |  | 8978 | 11021 |
| *Niwaella delicata* |  | 1020 |  |
| *Engraulis japonicus* |  | 33 |  |
| *Nipponocypris temminckii* |  |  | 7641 |
| *Oncorhynchus masou masou* |  |  | 2419 |
| *Silurus asotus* |  |  | 276 |
| *Homo sapiens* | 0 | 392 | 0 |

Table S5c Results of eDNA metabarcoding from the Yaizuchuo sample.

| *Scientific name* | reads (1) | reads (2) | reads (3) | reads (4) | reads (5) | reads (6) | reads (7) | reads (8) | reads (9) | reads (10) | reads (11) | reads (12) | reads (13) | reads (14) |
| --- | --- | --- | --- | --- | --- | --- | --- | --- | --- | --- | --- | --- | --- | --- |
| *Zacco platypus* | 7089 | 9730 | 11679 | 8188 | 8910 | 6211 | 8294 | 7029 | 6341 | 6646 | 8626 | 5901 | 6030 | 4599 |
| *Tridentiger* sp. | 2183 | 2420 | 5647 | 4658 | 1095 | 2193 | 959 | 2673 | 3453 | 4078 | 2315 | 1963 | 1652 | 1703 |
| *Rhinogobius* sp. | 1878 | 3990 | 5106 | 4178 | 2437 | 2593 | 2178 | 3527 | 2560 | 3602 | 2303 | 2452 | 1938 | 2391 |
| *Rhinogobius similis* | 871 | 654 | 2839 | 1717 | 1901 | 695 | 1284 | 943 | 1019 | 1516 | 1639 | 884 | 1639 | 667 |
| *Cobitis* sp. *BIWAE type B* | 550 | 596 | 1298 | 1014 | 246 | 359 | 224 | 481 | 394 | 393 | 364 | 251 | 321 | 334 |
| *Pseudaspius hakonensis* | 510 | 257 | 1238 | 961 | 431 | 446 | 294 | 552 | 860 | 378 | 332 |  | 109 |  |
| *Mugil cephalus* | 271 | 505 | 3381 | 2803 | 143 | 334 | 216 | 946 | 212 | 297 | 401 | 281 | 150 | 131 |
| *Pseudogobio esocinus* | 111 | 55 |  |  |  |  |  | 55 |  |  | 42 | 132 | 36 |  |
| *Cyprinus carpio* | 221 | 377 | 348 | 347 | 83 | 265 | 213 | 383 | 733 | 618 | 630 | 292 | 384 | 79 |
| *Carassius* sp. |  |  | 97 | 66 | 161 |  |  |  | 147 | 14 |  |  |  |  |
| *Sicyopterus japonicus* | 134 |  | 723 | 9699 | 156 | 189 | 98 | 3425 | 303 | 322 | 123 | 63 | 42 | 81 |
| *Gymnogobius petschiliensis* | 120 |  |  | 366 |  |  |  |  | 58 |  |  |  |  |  |
| *Anguilla japonica* | 86 |  |  |  | 52 | 42 |  | 127 | 53 | 44 |  |  | 50 |  |
| *Lepomis* sp. | 41 |  |  |  |  |  |  |  |  |  |  |  |  |  |
| *Pseudorasbora parva* |  |  |  |  |  |  |  |  | 33 |  |  |  |  |  |
| *Plecoglossus altivelis altivelis* |  |  | 303 | 185 | 62 | 43 | 35 | 105 |  |  |  | 31 |  |  |
| *Silurus* sp. | 295 |  |  |  |  | 18 |  |  |  | 41 |  |  |  | 16 |
| *Nipponocypris temminckii* |  |  | 207 | 71 |  |  |  |  |  |  |  |  |  |  |
| *Oreochromis* sp. |  |  |  |  |  | 42 |  |  |  |  |  |  |  |  |
| *Oreochromis niloticus* |  |  |  |  |  |  |  |  |  |  |  | 265 |  |  |
| *Micropterus salmoides salmoides* |  |  |  |  |  |  |  |  |  |  |  |  |  | 109 |
| *Acheilognathus rhombeus* |  |  |  |  |  |  |  |  |  |  |  |  |  |  |
| *Homo sapiens* |  | 73 | 2325 | 745 | 80 |  | 117 | 65 |  | 60 |  |  |  |  |
| *Sus scrofa* |  |  |  |  | 78 |  |  |  |  |  |  |  |  |  |

Numbers in cells indicate the number of reads obtained using the MitoFish pipeline. Reads (n) indicate the results obtained from each repeated sampl

Table S6a. Number of students who had an increase or decrease in responses regarding preferences for organisms through environmental education.

| Question | Comparison | Water sampling | Number of change responses. | *p* value  （χ^2^ test） |
| --- | --- | --- | --- | --- |
| Do you like biology? | QS1 vs. QS2 | + | up: 71, down: 37 | up: 0.07  down: 1.00 |
|  |  | - | up: 22, down: 18 |  |
|  | QS2 vs. QS3 | + | up: 62, down: 41 | up: 0.95  down: 0.06 |
|  |  | - | up: 28, down: 10 |  |
| Do you like organisms? | QS1 vs. QS2 | + | up: 51, down: 54 | up: 0.55  down: 0.87 |
|  |  | - | up: 20, down: 27 |  |
|  | QS2 vs. QS3 | + | up: 64, down: 48 | up: 0.83  down: 0.08 |
|  |  | - | up: 32, down: 13 |  |
| Are you willing to touch organisms? | QS1 vs. QS2 | + | up: 66, down: 62 | up: 0.21  down: 0.09 |
|  |  | - | up: 23, down: 19 |  |
|  | QS2 vs. QS3 | + | up: 78, down: 46 | up: 0.07  down: 0.57 |
|  |  | - | up: 25, down: 18 |  |

Table S6b. Number of students who had an increase or decrease in responses regarding biodiversity through environmental education.

| Question | Comparison | water Sampling | number of change responses. | *p* value  （χ^2^ test） |
| --- | --- | --- | --- | --- |
| The natural environment (mountains, rivers, sea, satoyama, rice paddies, nature parks, etc.) of my local area is well preserved. | QS1 vs. QS2 | + | up: 59, down: 68 | up: 0.34  down: 0.37 |
|  |  | - | up: 34, down: 26 |  |
|  | QS2 vs. QS3 | + | up: 85, down: 37 | up: 0.60  down: 0.30 |
|  |  | - | up: 44, down: 12 |  |
| The grounds of the XX High School (the name of the high school where the program was conducted) are rich in nature. | QS1 vs. QS2 | + | up: 65, down: 47 | up: 0.89  down: **< 0.05** |
|  |  | - | up: 32, down: 39 |  |
|  | QS2 vs. QS3 | + | up: 71, down: 42 | up: 0.92  down: 0.41 |
|  |  | - | up: 32, down: 15 |  |
| I support prioritizing disaster prevention measures such as riverbank protection work and mountain slope maintenance in my area, even if they negatively impact natural ecosystems. | QS1 vs. QS2 | + | up: 93, down: 51 | up: 1.0  down: 0.77 |
|  |  | - | up: 43, down: 30 |  |
|  | QS2 vs. QS3 | + | up: 93, down: 57 | up: 0.92  down: 0.79 |
|  |  | - | up: 41, down: 29 |  |
| I support using tax funds to preserve and enrich the diverse natural ecosystems in my local mountains and riparian areas. | QS1 vs. QS2 | + | up: 84, down: 42 | up: **< 0.05**  down: 0.28 |
|  |  | - | up: 25, down: 26 |  |
|  | QS2 vs. QS3 | + | up: 53, down: 55 | up: 0.03  down: **< 0,05** |
|  |  | - | up: 38, down: 11 |  |
| The natural environment where I live contains many different species. | QS1 vs. QS2 | + | up: 94, down: 47 | up: 0.26  down: 0.01 |
|  |  | - | up: 36, down: 37 |  |
|  | QS2 vs. QS3 | + | up: 75, down: 42 | up: 0.26  down: 0.30 |
|  |  | - | up: 43, down: 14 |  |
| Many different species inhabit the river near our high school. | QS1 vs. QS2 | + | up: 123,  down: 37 | up: **< 0.05**  down: 1.0 |
|  |  | - | up: 42, down: 18 |  |
|  | QS2 vs. QS3 | + | up: 91, down: 33 | up: 0.06  down: 0.86 |
|  |  | - | up: 56, down: 17 |  |
| I support removing organisms from my local area if they pose risks to humans. | QS1 vs. QS2 | + | up: 63, down: 68 | up: 0.89  down: 0.69 |
|  |  | - | up: 28, down: 35 |  |
|  | QS2 vs. QS3 | + | up: 59, down: 70 | up: 0.21  down: 0.72 |
|  |  | - | up: 20, down: 30 |  |
| I support removing organisms from my local area solely because of their strange appearance even if they cause no harm. | QS1 vs. QS2 | + | up: 41, down: 81 | up: 0.99  down: 0.50 |
|  |  | - | up: 20, down: 33 |  |
|  | QS2 vs. QS3 | + | up: 63, down: 66 | up: 0.05  down: 0.71 |
|  |  | - | up: 18, down: 28 |  |
| I support using tax funds to protect rare species in my local area. | QS1 vs. QS2 | + | up:70, down: 44 | up: 0.60  down: **< 0.05** |
|  |  | - | up: 29, down: 32 |  |
|  | QS2 vs. QS3 | + | up: 65, down: 51 | up: 0.43  down: 0.32 |
|  |  | - | up: 36, down: 18 |  |
| I support using tax funds to remove invasive species in my local area. | QS1 vs. QS2 | + | up: 67, down: 44 | up: 0.89  down: 0.38 |
|  |  | - | up: 33, down: 26 |  |
|  | QS2 vs. QS3 | + | up: 69, down: 61 | up: 0.43  down: 0.11 |
|  |  | - | up: 27, down: 19 |  |
| I support protecting the local ecosystems by donating money or participating in conservation activities. | QS1 vs. QS2 | + | up: 96, down: 24 | up: **< 0.05**  down: 0.15 |
|  |  | - | up: 28, down: 18 |  |
|  | QS2 vs. QS3 | + | up: 58, down: 50 | up: 0.38  down: 0.13 |
|  |  | - | up: 33, down: 15 |  |
| I support protecting biodiversity even if it requires sacrificing some modern conveniences (e.g. electricity usage and infrastructure development). | QS1 vs. QS2 | + | up: 89, down: 47 | up: 1.0  down: 0.18 |
|  |  | - | up: 41, down: 30 |  |
|  | QS2 vs. QS3 | + | up: 75, down: 63 | up: 0.33  down: 0.31 |
|  |  | - | up: 42, down: 23 |  |

Table S6c. Number of students who had an increase or decrease in responses regarding ecosystem service through environmental education.

| Question | Comparison | water Sampling | number of change responses. | *p* value  （χtest） |
| --- | --- | --- | --- | --- |
| Do you think that the ecosystem services in your local area should be improved? | QS1 vs. QS2 | + | up: 101,  down: 21 | up: 0.11  down: 0.91 |
|  |  | - | up: 36, down: 11 |  |
|  | QS2 vs. QS3 | + | up: 71, down: 40 | up: 0.93  down: 0.52 |
|  |  | - | up: 32, down: 15 |  |
| Do you think that the ecosystem disservices in your local area should be controlled? | QS1 vs. QS2 | + | up: 111,  down: 35 | up: 0.09  down: 1.00 |
|  |  | - | up: 40, down: 16 |  |
|  | QS2 vs. QS3 | + | up: 57, down: 68 | up: 1.00  down: 0.16 |
|  |  | - | up: 26, down: 23 |  |
| Do you think that an increase in biodiversity will also increase ecosystem services? | QS1 vs. QS2 | + | up: 92, down: 23 | up: 0.40  down: 0.53 |
|  |  | - | up: 37, down: 14 |  |
|  | QS2 vs. QS3 | + | up: 58, down: 47 | up: 0.24  down: 0.80 |
|  |  | - | up: 20, down: 20 |  |
| Do you think that an increase in biodiversity will also increase ecosystem disservices? | QS1 vs. QS2 | + | up: 89, down: 29 | up: **< 0.001**  down: **< 0.05** |
|  |  | - | up: 73, down: 24 |  |
|  | QS2 vs. QS3 | + | up: 54, down: 35 | up: 1.00  down: 0.54 |
|  |  | - | up: 25, down: 20 |  |

The water sampling column indicates whether the water was sampled in our program. + indicates that water sampling was performed, and - indicates that water sampling was not performed. The comparison column indicates which of the questionnaires (QS1, QS2, or QS3) were compared. The number of responses column shows the number of students whose responses increased or decreased between questionnaires. The *p* value column shows the results for students who collected water samples and those wh o did not.

Table S7 Results of logistic regression analysis of students who collected water samples.

| question | up/down | comparison | estimate_ex | estimate_like | estimate_touch | p.value_ex | p.value_like | p.value_touch |
| --- | --- | --- | --- | --- | --- | --- | --- | --- |
| The natural environment (mountains, rivers, sea, satoyama, rice paddies, nature parks, etc.) of my local area is well preserved. | up | QS1 vs. QS2 | 0.04 | -0.20 | 0.08 | 0.90 | 0.35 | 0.69 |
|  | up | QS2 vs. QS3 | 0.00 | 0.12 | 0.12 | 0.99 | 0.55 | 0.49 |
|  | down | QS1 vs. QS2 | 0.33 | 0.17 | 0.02 | 0.29 | 0.40 | 0.91 |
|  | down | QS2 vs. QS3 | -0.20 | -0.32 | 0.28 | 0.60 | 0.22 | 0.25 |
| The grounds of the XX High School (the name of the high school where the program was conducted) are rich in nature. | up | QS1 vs. QS2 | -0.14 | 0.30 | -0.25 | 0.63 | 0.15 | 0.19 |
|  | up | QS2 vs. QS3 | -0.14 | -0.25 | 0.27 | 0.63 | 0.21 | 0.15 |
|  | down | QS1 vs. QS2 | 0.25 | 0.32 | -0.02 | 0.49 | 0.19 | 0.94 |
|  | down | QS2 vs. QS3 | -0.23 | 0.01 | -0.03 | 0.51 | 0.98 | 0.88 |
| I support prioritizing disaster prevention measures such as riverbank protection work and mountain slope maintenance in my area, even if they negatively impact natural ecosystems. | up | QS1 vs. QS2 | -0.12 | 0.09 | 0.01 | 0.65 | 0.62 | 0.94 |
|  | up | QS2 vs. QS3 | -0.05 | 0.13 | -0.01 | 0.86 | 0.49 | 0.96 |
|  | down | QS1 vs. QS2 | -0.03 | -0.05 | -0.23 | 0.92 | 0.82 | 0.27 |
|  | down | QS2 vs. QS3 | -0.13 | -0.33 | 0.17 | 0.69 | 0.13 | 0.40 |
| I support using tax funds to preserve and enrich the diverse natural ecosystems in my local mountains and riparian areas. | up | QS1 vs. QS2 | -0.04 | -0.55 | 0.52 | 0.88 | **< 0.05** | **< 0.05** |
|  | up | QS2 vs. QS3 | 0.40 | -0.01 | -0.07 | 0.25 | 0.96 | 0.74 |
|  | down | QS1 vs. QS2 | -0.55 | 0.34 | 0.04 | 0.11 | 0.19 | 0.87 |
|  | down | QS2 vs. QS3 | -0.09 | -0.14 | 0.04 | 0.79 | 0.52 | 0.83 |
| The natural environment where I live contains many different species. | up | QS1 vs. QS2 | 0.15 | -0.07 | -0.24 | 0.59 | 0.70 | 0.15 |
|  | up | QS2 vs. QS3 | -0.18 | -0.20 | 0.39 | 0.54 | 0.32 | **< 0.05** |
|  | down | QS1 vs. QS2 | 0.27 | 0.24 | 0.19 | 0.46 | 0.33 | 0.37 |
|  | down | QS2 vs. QS3 | 0.03 | 0.05 | -0.01 | 0.94 | 0.84 | 0.96 |
| Many different species inhabit the river near our high school. | up | QS1 vs. QS2 | 0.09 | -0.16 | -0.02 | 0.74 | 0.36 | 0.90 |
|  | up | QS2 vs. QS3 | -0.02 | 0.19 | -0.10 | 0.93 | 0.30 | 0.55 |
|  | down | QS1 vs. QS2 | -0.19 | -0.30 | 0.29 | 0.61 | 0.25 | 0.23 |
|  | down | QS2 vs. QS3 | -0.54 | -0.25 | 0.24 | 0.15 | 0.37 | 0.35 |
| I support removing organisms from my local area if they pose risks to humans. | up | QS1 vs. QS2 | -0.37 | 0.00 | 0.13 | 0.22 | 0.99 | 0.48 |
|  | up | QS2 vs. QS3 | -0.03 | -0.51 | 0.56 | 0.93 | **< 0.05** | **< 0.05** |
|  | down | QS1 vs. QS2 | -0.20 | 0.22 | 0.08 | 0.49 | 0.30 | 0.66 |
|  | down | QS2 vs. QS3 | 0.64 | -0.32 | 0.02 | 0.05 | 0.11 | 0.93 |
| I support removing organisms from my local area solely because of their strange appearance even if they cause no harm. | up | QS1 vs. QS2 | -0.24 | 0.60 | -0.23 | 0.49 | **< 0.05** | 0.30 |
|  | up | QS2 vs. QS3 | 0.62 | -0.16 | 0.33 | 0.07 | 0.47 | 0.09 |
|  | down | QS1 vs. QS2 | 0.52 | 0.00 | -0.20 | 0.09 | 0.99 | 0.25 |
|  | down | QS2 vs. QS3 | -0.40 | 0.00 | -0.13 | 0.18 | 0.99 | 0.50 |
| I support using tax funds to protect rare species in my local area. | up | QS1 vs. QS2 | 0.00 | -0.10 | 0.08 | 0.99 | 0.60 | 0.67 |
|  | up | QS2 vs. QS3 | 0.04 | -0.22 | 0.18 | 0.91 | 0.29 | 0.34 |
|  | down | QS1 vs. QS2 | -0.49 | -0.33 | 0.36 | 0.15 | 0.18 | 0.11 |
|  | down | QS2 vs. QS3 | -0.21 | 0.11 | -0.08 | 0.51 | 0.63 | 0.69 |
| I support using tax funds to remove invasive species in my local area. | up | QS1 vs. QS2 | 0.49 | -0.31 | 0.30 | 0.13 | 0.14 | 0.12 |
|  | up | QS2 vs. QS3 | 0.21 | -0.38 | 0.05 | 0.50 | 0.06 | 0.79 |
|  | down | QS1 vs. QS2 | 0.36 | 0.15 | -0.21 | 0.34 | 0.53 | 0.33 |
|  | down | QS2 vs. QS3 | 0.11 | 0.15 | -0.15 | 0.73 | 0.49 | 0.42 |
| I support protecting the local ecosystems by donating money or participating in conservation activities. | up | QS1 vs. QS2 | -0.36 | -0.25 | 0.10 | 0.17 | 0.17 | 0.55 |
|  | up | QS2 vs. QS3 | 1.23 | 0.06 | -0.31 | **< 0.05** | 0.78 | 0.12 |
|  | down | QS1 vs. QS2 | 0.35 | -0.15 | 0.15 | 0.48 | 0.62 | 0.60 |
|  | down | QS2 vs. QS3 | -0.24 | -0.03 | 0.11 | 0.46 | 0.91 | 0.59 |
| I support protecting biodiversity even if it requires sacrificing some modern conveniences (e.g. electricity usage and infrastructure development). | up | QS1 vs. QS2 | -0.17 | -0.01 | -0.09 | 0.54 | 0.97 | 0.59 |
|  | up | QS2 vs. QS3 | -0.14 | -0.32 | 0.14 | 0.62 | 0.10 | 0.45 |
|  | down | QS1 vs. QS2 | -0.62 | -0.16 | -0.07 | 0.06 | 0.48 | 0.75 |
|  | down | QS2 vs. QS3 | -0.02 | -0.03 | 0.02 | 0.94 | 0.90 | 0.90 |
| Do you think that the ecosystem services in your local area should be improved? | up | QS1 vs. QS2 | 0.20 | -0.01 | 0.03 | 0.47 | 0.97 | 0.85 |
|  | up | QS2 vs. QS3 | -0.25 | -0.30 | 0.14 | 0.39 | 0.13 | 0.44 |
|  | down | QS1 vs. QS2 | -0.13 | 0.13 | -0.41 | 0.78 | 0.69 | 0.17 |
|  | down | QS2 vs. QS3 | 0.25 | 0.32 | -0.04 | 0.52 | 0.22 | 0.85 |
| Do you think that the ecosystem disservices in your local area should be controlled? | up | QS1 vs. QS2 | 0.15 | 0.05 | 0.12 | 0.58 | 0.77 | 0.44 |
|  | up | QS2 vs. QS3 | 0.31 | -0.46 | 0.41 | 0.35 | 0.04 | 0.05 |
|  | down | QS1 vs. QS2 | 0.31 | -0.05 | 0.01 | 0.45 | 0.84 | 0.97 |
|  | down | QS2 vs. QS3 | 0.64 | -0.04 | 0.22 | 0.05 | 0.87 | 0.26 |
| Do you think that an increase in biodiversity will also increase ecosystem services? | up | QS1 vs. QS2 | -0.11 | -0.02 | -0.19 | 0.69 | 0.90 | 0.26 |
|  | up | QS2 vs. QS3 | 0.35 | -0.13 | 0.20 | 0.30 | 0.56 | 0.31 |
|  | down | QS1 vs. QS2 | 0.53 | -0.26 | 0.16 | 0.31 | 0.41 | 0.59 |
|  | down | QS2 vs. QS3 | -0.36 | -0.04 | 0.07 | 0.27 | 0.87 | 0.76 |
| Do you think that an increase in biodiversity will also increase ecosystem disservices? | up | QS1 vs. QS2 | -0.17 | 0.17 | -0.24 | 0.54 | 0.34 | 0.16 |
|  | up | QS2 vs. QS3 | 0.01 | 0.02 | 0.09 | 0.98 | 0.93 | 0.65 |
|  | down | QS1 vs. QS2 | 0.23 | -0.09 | 0.24 | 0.61 | 0.77 | 0.37 |
|  | down | QS2 vs. QS3 | -0.13 | -0.03 | 0.18 | 0.73 | 0.91 | 0.47 |

Each column displays the following information: Question: The text of each question. up/down: indicates the dependent variable used, where up is a binary dataset where 1 is used for responses that increased between questionnaires, and 0 is used for responses that did not, and down is a binary dataset where 1 is used for responses that decreased between questionnaires, and 0 is used for responses that did not. Comparison: This indicates that the questionnaires were compared. estimate_ex, like, touch: These show results of estimates for the explanatory variables of past natural experiences of students, whether the students held a positive opinion of organisms, and whether they expressed desire to touch them. p.value_ex, like touch: This indicates the significance of the differences for each explanatory variable. Bold text indicates significant differences.

Table S8 Results of logistic regression analysis of students who did not collect water samples.

| question | up/down | comparison | estimate_ex | estimate_like | estimate_touch | *p* value_ex | *p* value_like | *p* value_touch |
| --- | --- | --- | --- | --- | --- | --- | --- | --- |
| The natural environment (mountains, rivers, sea, satoyama, rice paddies, nature parks, etc.) of my local area is well preserved. | up | QS1vs. QS2 | 0.08 | 0.02 | 0.07 | 0.87 | 0.95 | 0.79 |
|  | up | QS2 vs. QS3 | -0.11 | -0.10 | 0.20 | 0.81 | 0.73 | 0.41 |
|  | down | QS1vs. QS2 | -0.41 | 0.04 | -0.15 | 0.42 | 0.91 | 0.58 |
|  | down | QS2 vs. QS3 | 1.16 | 0.33 | -0.62 | 0.28 | 0.45 | 0.10 |
| The grounds of the XX High School (the name of the high school where the program was conducted) are rich in nature. | up | QS1vs. QS2 | 0.82 | 0.18 | -0.15 | 0.16 | 0.56 | 0.57 |
|  | up | QS2 vs. QS3 | 0.53 | 0.09 | 0.10 | 0.33 | 0.77 | 0.71 |
|  | down | QS1vs. QS2 | -0.17 | -0.03 | -0.09 | 0.71 | 0.92 | 0.71 |
|  | down | QS2 vs. QS3 | -0.02 | -0.46 | 0.04 | 0.97 | 0.23 | 0.90 |
| I support prioritizing disaster prevention measures such as riverbank protection work and mountain slope maintenance in my area, even if they negatively impact natural ecosystems. | up | QS1vs. QS2 | 0.17 | -0.03 | -0.35 | 0.72 | 0.91 | 0.15 |
|  | up | QS2 vs. QS3 | -0.68 | -0.19 | 0.09 | 0.12 | 0.50 | 0.73 |
|  | down | QS1vs. QS2 | -0.67 | -0.41 | 0.25 | 0.16 | 0.19 | 0.37 |
|  | down | QS2 vs. QS3 | 1.13 | 0.58 | -0.38 | 0.09 | 0.09 | 0.17 |
| I support using tax funds to preserve and enrich the diverse natural ecosystems in my local mountains and riparian areas. | up | QS1vs. QS2 | -0.27 | 0.05 | -0.41 | 0.62 | 0.89 | 0.14 |
|  | up | QS2 vs. QS3 | -0.57 | 0.15 | -0.06 | 0.20 | 0.61 | 0.82 |
|  | down | QS1vs. QS2 | -0.09 | -0.29 | 0.51 | 0.86 | 0.40 | 0.10 |
|  | down | QS2 vs. QS3 | -0.39 | 0.27 | -0.54 | 0.59 | 0.55 | 0.16 |
| The natural environment where I live contains many different species. | up | QS1vs. QS2 | 0.95 | -0.31 | 0.18 | 0.10 | 0.29 | 0.49 |
|  | up | QS2 vs. QS3 | 0.73 | 0.06 | 0.06 | 0.15 | 0.83 | 0.82 |
|  | down | QS1vs. QS2 | -0.03 | -0.13 | 0.12 | 0.94 | 0.64 | 0.63 |
|  | down | QS2 vs. QS3 | 0.06 | 0.59 | -0.47 | 0.93 | 0.19 | 0.18 |
| Many different species inhabit the river near our high school. | up | QS1vs. QS2 | -0.74 | 0.29 | -0.17 | 0.09 | 0.31 | 0.48 |
|  | up | QS2 vs. QS3 | 0.55 | 0.03 | -0.05 | 0.22 | 0.91 | 0.83 |
|  | down | QS1vs. QS2 | 0.31 | -0.27 | 0.15 | 0.65 | 0.46 | 0.66 |
|  | down | QS2 vs. QS3 | -0.48 | 0.11 | -0.04 | 0.41 | 0.78 | 0.90 |
| I support removing organisms from my local area if they pose risks to humans. | up | QS1vs. QS2 | -0.02 | -0.38 | 0.40 | 0.97 | 0.24 | 0.17 |
|  | up | QS2 vs. QS3 | -0.10 | 0.60 | -0.02 | 0.86 | 0.16 | 0.94 |
|  | down | QS1vs. QS2 | 0.12 | 0.08 | -0.04 | 0.81 | 0.79 | 0.86 |
|  | down | QS2 vs. QS3 | 0.40 | -0.13 | 0.25 | 0.46 | 0.68 | 0.39 |
| I support removing organisms from my local area solely because of their strange appearance even if they cause no harm. | up | QS1vs. QS2 | 0.12 | -0.05 | 0.21 | 0.84 | 0.89 | 0.53 |
|  | up | QS2 vs. QS3 | -0.04 | -0.09 | 0.24 | 0.95 | 0.81 | 0.49 |
|  | down | QS1vs. QS2 | 0.84 | 0.17 | -0.29 | 0.15 | 0.57 | 0.28 |
|  | down | QS2 vs. QS3 | 0.66 | 0.25 | -0.02 | 0.27 | 0.46 | 0.94 |
| I support using tax funds to protect rare species in my local area. | up | QS1vs. QS2 | -0.89 | 0.14 | -0.16 | 0.06 | 0.67 | 0.55 |
|  | up | QS2 vs. QS3 | 0.96 | -0.25 | 0.21 | 0.10 | 0.39 | 0.43 |
|  | down | QS1vs. QS2 | 0.76 | -0.19 | 0.09 | 0.19 | 0.52 | 0.74 |
|  | down | QS2 vs. QS3 | -0.08 | -0.43 | 0.41 | 0.89 | 0.25 | 0.25 |
| I support using tax funds to remove invasive species in my local area. | up | QS1vs. QS2 | 0.58 | 0.44 | -0.46 | 0.29 | 0.16 | 0.08 |
|  | up | QS2 vs. QS3 | 0.20 | -0.02 | -0.04 | 0.71 | 0.96 | 0.89 |
|  | down | QS1vs. QS2 | -0.11 | -0.17 | 0.31 | 0.84 | 0.61 | 0.31 |
|  | down | QS2 vs. QS3 | -0.42 | -0.20 | -0.18 | 0.46 | 0.56 | 0.57 |
| I support protecting the local ecosystems by donating money or participating in conservation activities. | up | QS1vs. QS2 | 0.22 | 0.07 | -0.36 | 0.68 | 0.83 | 0.19 |
|  | up | QS2 vs. QS3 | -1.15 | -0.46 | 0.26 | **< 0.05** | 0.13 | 0.35 |
|  | down | QS1vs. QS2 | -1.42 | -0.60 | 0.36 | **< 0.05** | 0.11 | 0.30 |
|  | down | QS2 vs. QS3 | 0.54 | -0.40 | 0.07 | 0.50 | 0.31 | 0.84 |
| I support protecting biodiversity even if it requires sacrificing some modern conveniences (e.g. electricity usage and infrastructure development). | up | QS1vs. QS2 | 0.96 | 0.36 | -0.16 | 0.07 | 0.22 | 0.52 |
|  | up | QS2 vs. QS3 | -0.64 | -0.33 | 0.23 | 0.15 | 0.24 | 0.37 |
|  | down | QS1vs. QS2 | -0.38 | -0.10 | 0.17 | 0.43 | 0.74 | 0.54 |
|  | down | QS2 vs. QS3 | 1.28 | 0.35 | -0.05 | 0.10 | 0.36 | 0.88 |
| Do you think that the ecosystem services in your local area should be improved? | up | QS1vs. QS2 | 0.11 | 0.03 | -0.27 | 0.82 | 0.91 | 0.28 |
|  | up | QS2 vs. QS3 | 0.79 | -0.42 | 0.41 | 0.18 | 0.18 | 0.15 |
|  | down | QS1vs. QS2 | 0.22 | 0.27 | -0.37 | 0.79 | 0.56 | 0.34 |
|  | down | QS2 vs. QS3 | 1.38 | -0.34 | 0.11 | 0.19 | 0.40 | 0.78 |
| Do you think that the ecosystem disservices in your local area should be controlled? | up | QS1vs. QS2 | -0.51 | -0.29 | 0.35 | 0.26 | 0.32 | 0.18 |
|  | up | QS2 vs. QS3 | 0.44 | -0.12 | -0.03 | 0.45 | 0.70 | 0.92 |
|  | down | QS1vs. QS2 | -0.23 | -0.41 | 0.52 | 0.72 | 0.30 | 0.17 |
|  | down | QS2 vs. QS3 | -0.02 | -0.38 | 0.49 | 0.97 | 0.27 | 0.14 |
| Do you think that an increase in biodiversity will also increase ecosystem services? | up | QS1vs. QS2 | 0.45 | -0.34 | 0.40 | 0.38 | 0.25 | 0.14 |
|  | up | QS2 vs. QS3 | 0.54 | 0.32 | -0.21 | 0.42 | 0.40 | 0.51 |
|  | down | QS1vs. QS2 | -0.31 | 0.75 | -0.30 | 0.63 | 0.13 | 0.41 |
|  | down | QS2 vs. QS3 | -0.24 | -0.88 | 1.01 | 0.68 | **< 0.05** | **< 0.05** |
| Do you think that an increase in biodiversity will also increase ecosystem disservices? | up | QS1vs. QS2 | 0.35 | 0.96 | -5.12 | 0.67 | 0.10 | **< 0.001** |
|  | up | QS2 vs. QS3 | 1.35 | 0.21 | -0.09 | 0.08 | 0.56 | 0.77 |
|  | down | QS1vs. QS2 | 0.86 | 1.89 | 2.52 | 0.29 | **< 0.05** | **< 0.001** |
|  | down | QS2 vs. QS3 | -0.26 | -0.47 | 0.48 | 0.65 | 0.20 | 0.16 |

Each column displays the following information: Question: The text of each question. up/down: indicates the dependent variable used, where up is a binary dataset where 1 is used for responses that increased between questionnaires, and 0 is used for responses that did not, and down is a binary dataset where 1 is used for responses that decreased between questionnaires, and 0 is used for responses that did not. Comparison: This indicates that the questionnaires have been compared. estimate_ex, like, touch: These show results of estimates for explanatory variables of past natural experiences of students, whether the students held a positive opinion of organisms, and whether they wanted to touch them. *p* value_ex, like touch: This shows the significance of the difference in each explanatory variable. Bold text indicates significant differences.
