## Supplementary text for "Feasibility of an eDNA-based educational program for high school students"

S1. Water sampling by students

The students visited rivers near their schools and collected water samples. The students rom Hidakamioka, Ena, and Yaizuchuo high schools collected water samples from the Takahara, Agi, and Seto Rivers, respectively. Students collected surface water from rivers using disposable plastic cups and pooled the water in buckets. Pooled water (1 L) was collected in a plastic bottle, and 10% benzalkonium chloride (1 mL) was added to prevent DNA degradation (Yamanaka et al., 2017). Water was collected by students who choose any location within the designated area of the river. The number of replicates of the buckets depended on the number of students at each high school, and students from Hidakamioka, Ena, and Yaizuchuo high schools collected 2, 3, and 14 water replicates, respectively. All samples were refrigerated immediately after collection and transported to the laboratory. A 1-L bottle of deionized water was prepared for the field blank on each collection day. To minimize contamination, bottles and buckets were pre-bleached, and disposable plastic cups were rinsed with water from the sampling site before use.

S2. Sample filtration and DNA extraction

Samples were filtered within 2 d of collection. GF/F glass fiber filters (Whatman, 0.47-mm pore size) were used to filter 1 L of water. After filtration, the filters were wrapped in aluminum foil and stored in the freezer (-20 °C) until used for DNA extraction. A DNeasy Blood & Tissue Kit (QIAGEN) was used for DNA extraction. The DNA extraction protocol was mainly based on the manual published by the eDNA Society (Minamoto et al., 2021), with some modifications based on Wu and Minamoto (2023). After placing the filter in a Salivette (Sarstat), ATL buffer (400 µL) and Proteinase K solution (40 µL) were added, and the mixture was incubated at 56 °C for 30 min in an incubator. The mixture was then centrifuged at 3000 *x* g for 3 min. TE buffer (220 µL; pH 8.0) was added and the mixture was centrifuged at 3000 *x* g for 3 min. Buffer AL (400 µL) and 100% EtOH (400 µL) were added to the liquid collected at the bottom of the salivette and mix thoroughly by pipetting. Up to 650 µL of this mixture was transferred to a DNeasy Mini Spin Column and centrifuged at 6000 *x* g for 1 min, and the filter was then discarded. This procedure was repeated until all the mixture was consumed. The procedure corresponded to the protocol recommended by the manufacturer. Finally, approximately 100 µL of eDNA solution was recovered. DNA was stored in the freezer (-20 °C) until use.

S3. Library preparation and next-generation sequencing

To perform eDNA metabarcoding analysis of fish, a MiFish-U primer designed for the 12rRNA gene region of mitochondrial DNA was used in the first polymerase chain reaction (PCR) (Miya et al., 2015). The first PCR reaction mixture was prepared by mixing 1×KAPA HiFi HotStartReadyMix (KAPA Biosystems), each primer at a final concentration of 300 nM, and distilled water (DW) to make a total volume of 11 µL/reaction, and then 1 µL of extracted DNA template was added (note: the total volume was 12 µL/reaction). The first PCR was performed in quadruplicate per sample. PCR conditions were as follows 95 ℃ for 3 min at the beginning, followed by 40 cycles of 98 ℃ for 20 s, 65 ℃ for 15 s, and 72 ℃ for 15 s, and finally 72 ℃ for 5 min. The products obtained after the first PCR were subjected to electrophoresis and checked to determine if the desired band length was amplified. If the desired band amplification could not be confirmed, the first PCR was repeated using KOD-Plus-Neo (TOYOBO) or Platinum SuperFi Ⅱ DNA Polymerase (Thermo Fisher Scientific) as the PCR enzyme. When KOD-Plus-Neo was used, the reaction mixture was prepared by mixing 0.25 U KOD-Plus-Neo polymerase, 1 × buffer, approximately 200 nM dNTP, approximately 1.56 nM MgSO_4_, and 625 nM of each primer (i.e., final concentration). DW was then added to a total volume of 10 µL/reaction, and 2 µL of extracted DNA template was then added (note: the total volume was 12 µL/reaction). PCR was performed in quadruplicate per sample. PCR conditions were as follows 94 ℃ for 2 min at the start, followed by a cycle of 98 °C for 10 s, 65 ℃ for 30 s, and 68 ℃ for 25 s for 40 cycles, and finally 5 min at 68 ℃. When using Platinum SuperFi II DNA Polymerase, 1 × Platinum SuperFi II DNA Polymerase, 1 × buffer, approximately 160 nM dNTP mix, and 500 nM of each primer (i.e., final concentration) were combined. DW was then added to a total volume of 10 µL/reaction, and 2 µL of extracted DNA template was then added (note: total volume was 12 µL/reaction). PCR was performed quadruplicate for each sample. PCR conditions were as follows: 95 ℃ for 3 min at the beginning, followed by 38 cycles of 98 ℃ for 20 s, 65 ℃ for 15 s, and 72 ℃ for 15 s, and finally 72 ℃ for 5 min. For the initial PCR, a blank sample was prepared for each PCR experiment.

The product, in which the target band was confirmed by electrophoresis, was purified using SPRIselect (Beckman Coulter). Briefly, SPRIselect (45 µL) was added to the PCR product, vortexed, and allowed to stand for 1 min. The tube was then placed on a magnetic plate and the solution was collected and discarded. Then, 85% EtOH (180 µL) was added, and after standing for 30 s, the solution was collected and discarded. The tube was removed from the magnetic plate, 40 µL of TE buffer (pH 8.0) was added followed by vortexing, and the tube was then allowed to stand for 1 min. DNA was purified by placing the tube on a magnetic plate and the solution was collected. The purified DNA concentration was measured using a Qubit 3.0 Fluorometer (Invitrogen). To ensure that the amount of product after the second PCR was consistent between samples, the DNA concentration of each sample was diluted to 0.1 ng/µL using TE buffer (pH 8.0).

A second round of PCR was performed to add tags to distinguish between samples. The second PCR reaction mixture was prepared by mixing 1 × KAPA HiFi HotStartReadyMix and each tag primer at a final concentration of 300 nM, adding DW to make a total volume of 11 µL/reaction, and then adding 1 µL of extracted DNA template (note: total volume of 12 µL/reaction). PCR conditions were as follows: 95 ℃ for 3 min at the beginning, followed by 12 cycles of 98 ℃ for 20 s, 72 ℃ for 20 s, and finally 72 ℃ for 5 min, and finally 72 ℃ for 5 min.

After the second PCR, the product was extracted using 2% E-Gel SizeSelect II agarose gels (Thermo Fisher Scientific) to obtain the target length band. The target band was checked using a bioanalyzer (Agilent Technologies) to confirm the successful recovery. Next-generation sequencing was performed using an iSeq100 (illumina). The DNA concentration of samples after the BioAnalyzer was measured using a Qubit 3.0 Fluorometer, and samples were diluted with DW to 1.0 nM to prepare the library. To improve the quality of next-generation sequencing, PhiX (Illumina) was added at a ratio of 20% (PhiX) to 80% (library). The mixture (20 µL) was sequenced using an iSeq100 sequencer (Illumina). Upon completion of the next-generation sequencing run, the cartridge was immediately removed and discarded, and the generated data were collected.

S4. Bioinformatics

The next-generation sequencing data were processed using the MitoFish pipeline (Sato et al., 2018; Zhu et al., 2023). The most recent references were used at the time of analysis (v3.91 for Hidakamioka and Ena; v3.94 for Yaizuchuo), and zOTUs with a match rate of ≤ 98.5% were excluded. Sequences obtained from MitoFish were checked using BLAST to identify other species with similar match rates. If the MitoFish annotation assigned a species that was unlikely to have been present at the sampling site, it was reassigned to a species that was present at the sampling site among the references with the highest match rate in BLAST. In cases where there were multiple references with the highest match rate in BLAST and it was not possible to determine which species were present at the sampling site, the genus was assigned.

Several fish species were detected in the Yaizuchuo field and the PCR blanks. When these species were detected in the corresponding water samples, the number of reads in the blank was subtracted from that of the corresponding water sample (Sakata et al., 2021). One fish species, Kanehira (*Acheilognathus rhombeus*), was detected in 37 reads from 1 of the PCR blanks, but not in the corresponding water sample. We determined that the experiment was conducted in an environment where external contamination of approximately 37 reads could occur and removed fish species with ≤ 37 reads from these samples.

S5. eDNA metabarcoding analysis results

The number of raw reads was 878,461 (n = 30:19 water samples, five field blanks, and six PCR blanks), and the number of reads remaining after the MitoFish pipeline was 497,415.

Fish metabarcoding analysis was performed on samples from Hidakamioka. A total of 9 fish species were detected in Takahara River samples (Tables S4 and S5a). The MitoFish pipeline results indicated that the species was Formosan landlocked salmon (*Oncorhynchus masou formosanus*); however, we determined that this sequence was derived from landlocked salmon (*Oncorhynchus masou*) based on additional BLAST confirmation. In addition, the species identified as torrent sculpin (*Cottus rhotheus*) in the MitoFish pipeline results was identified as the Japanese fluvial sculpin (*Cottus pollux*) in the same manner.

As a result of performing fish metabarcoding analysis of samples from Ena, 16 fish taxa (including 3 that could only be identified to the genus level) were detected in samples collected from the Agi River (Tables S4 and S5b). In the MitoFish pipeline, sequences thought to belong to the carp genus (*Cyprinus* sp.) were determined to be common carp (*Cyprinus carpio*) based on blast confirmation.

As a result of the fish metabarcoding analysis of the samples from Yaizuchuo, 22 fish taxa (including 6 that could only be identified to the genus level) were detected in samples collected from the Seto River (Tables S4 and S5c). In the MitoFish pipeline, the sequences thought to be from the carp genus (*Cyprinus* sp.) were determined to be common carp (*C. carpio*) based on methods similar to those used to analyze the results from Ena High School.

In samples from Hidakamioka, Ena, and Yaizuchuo, 9, 16, and 22 fish taxa were detected, respectively.
